# Exploration-exploitation mechanisms in recurrent neural networks and human learners in restless bandit problems

**DOI:** 10.1101/2023.04.27.538570

**Authors:** D. Tuzsus, A. Brands, I. Pappas, J. Peters

## Abstract

A key feature of animal and human decision-making is to balance the exploration of unknown options for information gain (directed exploration) versus selecting known options for immediate reward (exploitation), which is often examined using restless bandit tasks. Recurrent neural network models (RNNs) have recently gained traction in both human and systems neuroscience work on reinforcement learning, due to their ability to show meta-learning of task domains. Here we comprehensively compared the performance of a range of RNN architectures as well as human learners on restless four-armed bandit problems. The best-performing architecture (LSTM network with computation noise) exhibited human-level performance. Computational modeling of behavior first revealed that both human and RNN behavioral data contain signatures of higher-order perseveration, i.e., perseveration beyond the last trial, but this effect was more pronounced in RNNs. In contrast, human learners, but not RNNs, exhibited a positive effect of uncertainty on choice probability (directed exploration). RNN hidden unit dynamics revealed that exploratory choices were associated with a disruption of choice predictive signals during states of low state value, resembling a win-stay-loose-shift strategy, and resonating with previous single unit recording findings in monkey prefrontal cortex. Our results highlight both similarities and differences between exploration behavior as it emerges in meta-learning RNNs, and computational mechanisms identified in cognitive and systems neuroscience work.

## Introduction

Reinforcement learning (RL) theory (Sutton & Barto, 2018) is of central importance in psychology, neuroscience, computational psychiatry and artificial intelligence as it accounts for how artificial and biological agents learn from reward and punishment. According to both the law of effect in psychology (Thorndike, 1927) and the reward hypothesis in machine learning (Sutton & Barto, 2018), agents optimize behavior to maximize reward and minimize punishment. In computational psychiatry, RL theory has yielded valuable insights into changes in learning and decision-making associated with different mental disorders (Huys et al., 2016; Maia & Frank, 2011; Yahata et al., 2017).

To maximize reward, agents have to solve the exploration-exploitation dilemma (Sutton & Barto, 2018) that can be stated as follows: Should one pursue actions that led to reward in the past (exploitation) or should one explore novel courses of action for information gain (exploration)? In stable environments, where action-outcome contingencies are stable over time, the exploration-exploitation dilemma can be effectively solved by first exploring all available actions to identify the most rewarding one, and subsequently exploiting this action. In contrast, in volatile environments, action-outcome contingencies change over time, such that exploration and exploitation need to be continuously balanced by an agent. A high level of exploitation would make an agent unable to adapt to changes in the environment, whereas an excess of exploration would reduce reward accumulation, as optimal actions would oftentimes not be selected.

A number of computational strategies have been proposed to address the exploration-exploitation tradeoff (Sutton & Barto, 2018). In *ε*-greedy and softmax choice rules, exploration is achieved via choice randomization. While such “random” exploration appears to be one core component of both human and animal exploration (Daw et al., 2006; Ebitz et al., 2018; Schulz & Gershman, 2019; Wilson et al., 2014, 2021), computational modeling of behavior strongly suggests that humans additionally use “directed” or strategic exploration strategies (Chakroun et al., 2020; Schulz & Gershman, 2019; Speekenbrink & Konstantinidis, 2015; Wiehler et al., 2021; Wilson et al., 2014, 2021). This is typically modeled via an “exploration bonus” parameter that increases the value of options with greater information value (Chakroun et al., 2020; Speekenbrink & Konstantinidis, 2015; Wiehler et al., 2021; Wu et al., 2018). In volatile environments, the uncertainty associated with the outcome of a specific action is often taken as a proxy for information gain (Wilson et al., 2021). Exploring uncertain courses of action can thus increase information gain, over and above a simpler random exploration strategy.

A further related process pervasive in human and animal behavior is perseveration, the tendency of an agent to repeat previous choices regardless of obtained reward. Perseveration can be categorized into first-order or higher-order perseveration. First-order perseveration refers to the tendency to repeat the choice of the previous trial (*choice*_*t*−1_), but in the higher-order case, perseveration can extend to choices n-trials back (*choice*_*t*−*n*_) (Lau & Glimcher, 2005). First-order perseveration in RL tasks was observed in rats (Ito & Doya, 2009), monkeys (Balcarras et al., 2016; Lau & Glimcher, 2005) and humans (Chakroun et al., 2020; Wiehler et al., 2021), and there is evidence for higher-order perseveration in monkeys (Lau & Glimcher, 2005) and humans (Gershman, 2020; Palminteri, 2023; Seymour et al., 2012). Recently, Palminteri (Palminteri, 2023) showed that accounting for higher-order perseveration in RL models of behavior improves the interpretability of other model parameters.

Note that perseveration and exploration are associated with opposite choice patterns. Whereas high perseveration implies “sticky” behavior repeating previous choices, exploration entails a higher proportion of switch trials. Notably, explicitly accounting for perseveration in RL models improves the sensitivity to detect directed exploration effects. High perseveration can attenuate estimates of directed exploration driving the estimates to be lower or even negative if perseveration is not accounted for in the computational model of behavior (Badre et al., 2012; Chakroun et al., 2020; Daw et al., 2006; Payzan-LeNestour, 2012; Wiehler et al., 2021; Worthy et al., 2013). Chakroun et al. (2020) showed that including a first-order perseveration term in the RL model increases estimates of directed exploration and improves model fit to human data. Thus, taking into account perseveration behavior is crucial when examining exploration.

In humans, exploratory choices are associated with increased activity in the fronto-parietal network (Beharelle et al., 2015; Chakroun et al., 2020; Daw et al., 2006; Wiehler et al., 2021) and regulated by dopamine and norepinephrine neuromodulatory systems (Chakroun et al., 2020; Cremer et al., 2023; Dubois et al., 2021; McClure et al., 2005; Swanson et al., 2020). Choice predictive signals in prefrontal cortex neural populations are disrupted during exploratory choices, reflecting a potential neural mechanism for random exploration (Ebitz et al., 2018).

Such neuroscientific lines of work have increasingly been informed by computational neuroscience approaches (Mante et al., 2013). Here, neural network models are applied to clarify the computational principles underlying task performance. In the context of RL problems, recurrent neural network models (RNNs) are particularly powerful tools. They constitute deep artificial neural network models for sequential data (LeCun et al., 2015) such as RL tasks (Botvinick et al., 2020). Agents interact with the environment, and receive environmental feedback (e.g., rewards), which then informs subsequent choices. RNNs can be applied to RL problems due to their recurrent connectivity pattern. Each time step, RNN hidden units receive information regarding the network’s activation state at the previous time step via recurrent connections, thereby endowing the network with memory about what has happened before. Training and analysis of such models offer potential novel insights with implications for neuroscience (Botvinick et al., 2020). For example, the representations that emerge in a network’s hidden unit activation pattern following training (or over the course of training) can be directly analyzed (Findling & Wyart, 2020; Mante et al., 2013; Tsuda et al., 2020; Wang et al., 2018), similar to the analysis of high-dimensional neural data (Cunningham & Yu, 2014; Ebitz et al., 2018; Mante et al., 2013). This can reveal insights into the computations and representations underlying a network’s performance.

Neural network modeling approaches can also complement computational modeling of behavior as typically done in psychology and cognitive neuroscience (Farrell & Lewandowsky, 2018; Wilson & Collins, 2019). Traditional computational modeling of behavior can be characterized as a theory-driven approach, where computational mechanisms and representations hypothesized to underlie performance of a given task are explicitly and rigidly build into a quantitative model. While this approach is helpful to compare candidate models, the rigid dependency of these models on built-in a priori assumptions preclude the discovery of novel mechanisms and representations that could underlie task performance. In contrast, neural network modeling can be characterized as a data-driven approach, where highly flexible neural networks are trained to solve specific tasks or problems. RNN dynamics and representations might then reveal novel potential mechanisms and representations that support similar tasks by virtue of the RNNs independent data-driven learning capacity (Botvinick et al., 2020). Reward learning (Findling & Wyart, 2020; Tsuda et al., 2020; Wang et al., 2018) and decision-making (Findling & Wyart, 2020; Mante et al., 2013) are prominent recent examples.

In this line of work, RNNs are trained to solve reinforcement learning and decision-making tasks from the human and animal neuroscience literature, and the mechanisms underlying their performance are examined. Note that this is distinct from using RNNs to model human behavioral data, where standard computational models are replaced by RNNs (Dezfouli et al., 2019). RNNs trained on such tasks achieved a form of “meta-learning”: when weights were held fixed following training, the models had acquired the ability to solve novel instantiations of tasks from the same task family (Dasgupta et al., 2019; Findling & Wyart, 2021; Tsuda et al., 2020; Wang et al., 2018). Reinforcement learning over a large number of training episodes via slow adjustments of network weights, gave rise to a much faster reinforcement learning algorithm embedded in the network dynamics, and not involving further weight changes (Botvinick et al., 2020; Findling & Wyart, 2021; Wang et al., 2018).

Finally, RNNs with noisy computations might be more resilient to adverse conditions (e.g. contingency reversals, volatility) than their counterparts with deterministic computations (Findling & Wyart, 2020). This resonates with findings from the machine learning literature suggesting improved performance of neural networks with noisy computations under some conditions (Dong et al., 2020; Fortunato et al., 2019; Qin & Vucinic, 2018). Likewise, mental representations (Drugowitsch et al., 2016) and neural representations (Findling et al., 2019; Findling & Wyart, 2021; Renart & Machens, 2014) might benefit from some degree of representational imprecision (e.g., representations infused with task-independent noise).

RNNs trained on cognitive tasks from the neuroscience literature can reveal how artificial neural networks diverge from (or converge with) human and animal behavior. Also, RNNs might show human-like behavior by mere statistical learning without the use of human-like abstract rule learning (Kumar et al., 2022). While deep RL agents show superhuman ability in games like Go, Shogi, Chess and Atari games (Mnih et al., 2015; Silver et al., 2017, 2018) they fail to perform better than chance level on a standard T-maze task from animal learning (Wauthier et al., 2021). One of the most prominent differences between human learners and neural networks is the number of interactions with the environment required to learn a task (Botvinick et al., 2019; Lake et al., 2015; Marcus, 2018; Tsividis et al., 2021). This is in part related to exploration inefficiency (“sampling inefficiency”) during training (Hao et al., 2023). Even though there is ongoing work on endowing deep RL agents with improved exploration strategies (Hao et al., 2023; Ladosz et al., 2022; Tsividis et al., 2021), there is so far limited evidence with respect to exploration strategies emerging in meta-learning RNNs. Binz & Schulz (Binz & Schulz, 2022) showed that the large language model (LLM) GPT-3 shows no evidence of directed exploration in the “Horizon Task” from human cognitive neuroscience (Wilson et al., 2014). However, LLMs are trained passively on vast amounts of text data, whereas humans learn actively by interacting with dynamic environments. Whether meta-learning RNNs trained on dynamic environments employ directed exploration is an open question.

Bandit tasks constitute a classical testing bed for RL agents (Sutton & Barto, 2018), and are regularly applied to study human and animal exploration (Beharelle et al., 2015; Chakroun et al., 2020; Daw et al., 2006; Ebitz et al., 2018; Findling et al., 2019, 2019; Hamid et al., 2016; Mohebi et al., 2019; Wiehler et al., 2021). In non-stationary (*restless*) bandit tasks, agents select among a number of options (“bandits”) with dynamically changing reinforcement rates or magnitudes. In contrast, in stationary bandit problems reinforcement rates are fixed. RNNs achieve state-of-the-art performance on stationary bandit tasks (Wang et al., 2018) and in reversal schedules (Behrens et al., 2007) adapt their learning rates to environmental volatility (Wang et al., 2018). Furthermore, RNNs with computation noise can solve restless bandit tasks when trained on stationary bandits (Findling & Wyart, 2020), in contrast to their counterparts with deterministic computations. Human exploration behavior in restless bandit tasks is typically better accounted for by models with dynamic uncertainty-dependent learning rates such as the Kalman Filter (Daw et al., 2006; Kalman, 1960). Furthermore, humans regularly apply a directed exploration strategy on restless bandit tasks. This is modeled using an additional “exploration bonus” parameter that typically takes on positive values, reflecting directed exploration of uncertain options (Beharelle et al., 2015; Chakroun et al., 2020; Speekenbrink & Konstantinidis, 2015; Wiehler et al., 2021; Wilson et al., 2021; Wu et al., 2018).

Initial work on RNN mechanisms supporting bandit task performance (Findling & Wyart, 2020; Song et al., 2017; Wang et al., 2018), have predominantly focused on stationary bandits (Wang et al., 2018). However, stationary bandits preclude a comprehensive analysis of exploration mechanisms, because exploration is restricted to the first few trials. Furthermore, previous work often focused on two-armed bandit problems (Findling et al., 2019; Findling & Wyart, 2020; Song et al., 2017). However, these tasks are limited in that only one alternative can be explored at any given point in time. Although previous work has begun to use classical computational modeling to better understand RNN behavior (Fintz et al., 2022; Wang et al., 2018), a comprehensive comparison of human and RNN behavior and computational mechanisms when solving the exact same RL problems is still lacking. In addition, similar to so-called researcher’s degrees of freedom in experimental work (Wicherts et al., 2016), the study of RNNs is associated with a large number of design choices, e.g. with respect to the specifics of the architecture, details of the training schemes as well as hyperparameter settings. Yet, a comprehensive comparison of different network architectures and design choices in the context of RL tasks from the human cognitive neuroscience literature is still lacking.

Here, we addressed these issues in the following ways. First, we comprehensively compared a large set of RNN architectures in terms of their ability to exhibit human-level performance on restless four-armed bandit problems (See Appendix). Note that RNNs were trained as stand-alone RL agents, and were not used to replace standard computational models to account for human behavior. Second, to compare computational strategies between RNNs and human subjects, we used comprehensive computationalmodeling of human and RNN behavior during performance of the exact same RL problems. Finally, we expanded upon previous approaches to the analysis of RNN hidden unit activity patterns (Findling & Wyart, 2020; Mante et al., 2013; Wang et al., 2018) by leveraging analysis approaches from systems neuroscience studies of exploration (Ebitz et al., 2018).

## Methods

We trained artificial recurrent neural networks (RNN) in a Meta-Reinforcement Learning framework, where the RNN agent is trained on a task family to enable quick adaptations to novel but related tasks without new training. This process can be characterized by two loops. During training an outer loop trains the RNN parameters and samples a task instance from a task distribution for each training episode. Within the inner loop the agent performs the sampled task instance. During each cycle of the outer loop the RNN parameters are improved to increase performance in the inner loop (See Fig. 1a). During test, training is completed and parameters of the RNN agent are held fixed. Testing now resolves only in the inner loop where the agent performs the restless bandit task instances in (Chakroun et al., 2020) without any new parameter updates, just by using the learned inner dynamics during training (See Fig. 1 b). Further. we comprehensively compared a large set of RNN architectures in terms of their ability to exhibit human-level performance on restless four-armed bandit problems varying various architectural design choices like cell type, noise, loss function and entropy regularization (See Fig. 2). For details regarding RNN architectures, training and testing procedure see Appendix.

**Fig. 1.**
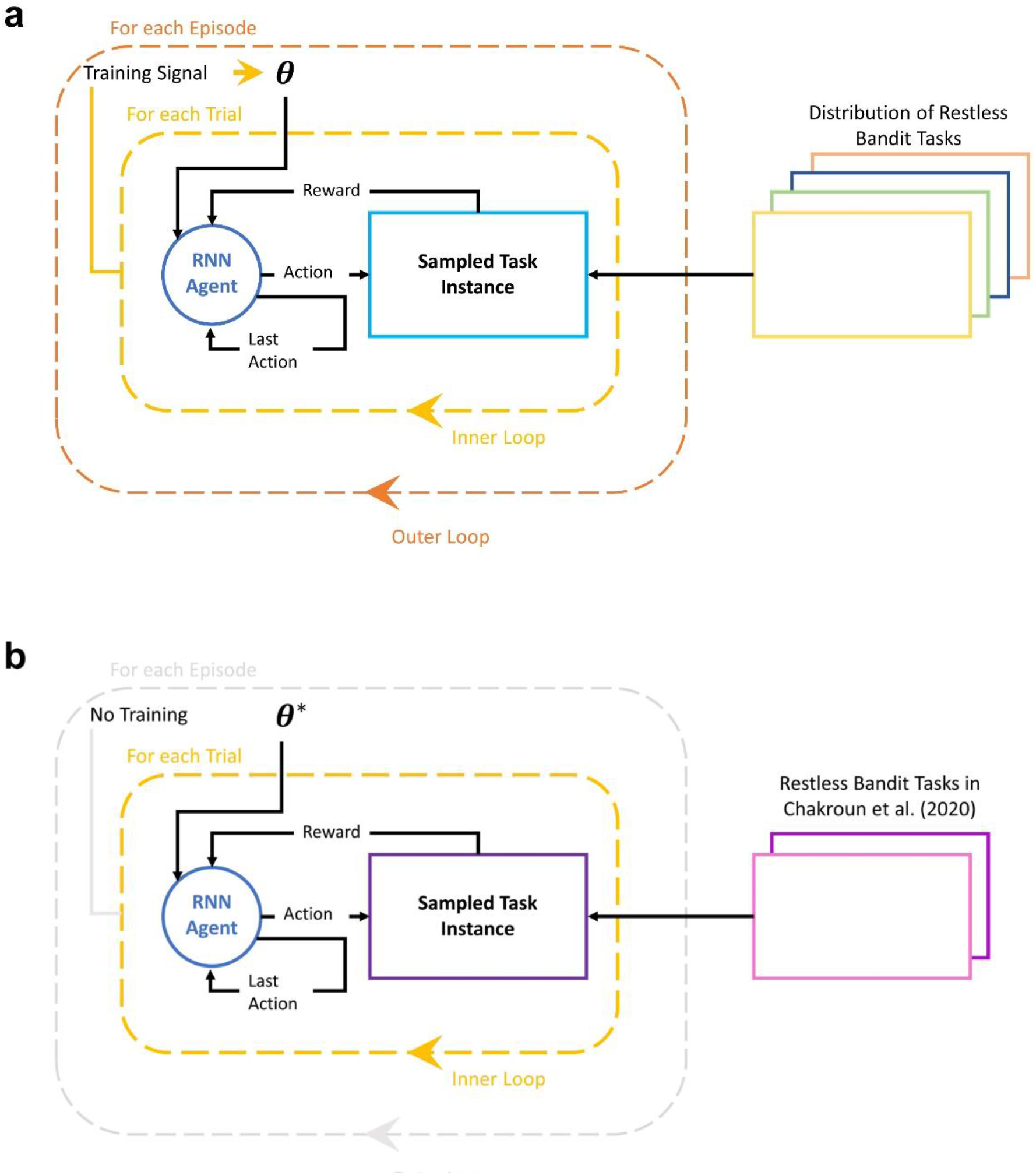
Schematic of Meta-Reinforcement Learning illustrated as Inner and Outer Loops. (a) Training loop: For each episode the outer loop trains the RNN parameters θ (e.g., weights and biases), which constitutes the agent in the inner loop. The inner loop iterates over trials of the sampled task instance, where the agent performs the task. In the outer loop, for every cycle, a new task instance is sampled from a distribution of restless bandit tasks with common structure. (b) Testing loop: After training completion, the outer loop is omitted and trained RNN parameters **θ**^**\***^ are held fixed. Testing resolves in the inner loop, where the agent performs the restless bandit tasks in Chakroun et al. (2020) without any new parameter updates, just by using the learned inner dynamics during training. Figures adapted and modified from Botvinick et al. (2019).

**Fig. 2.**
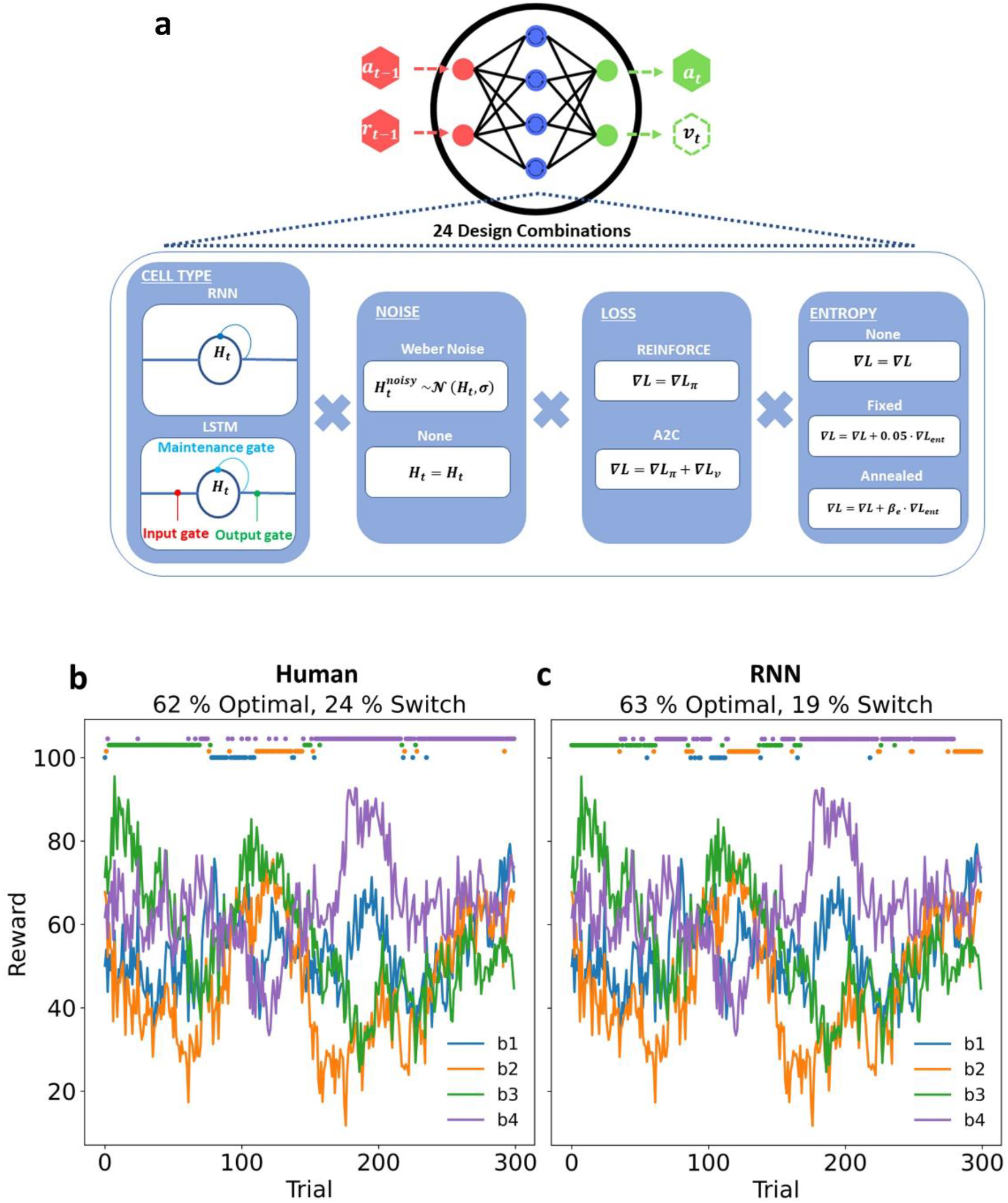
Artificial agent architectures and comparison of task performance to human agent. (a) The input to the artificial agent is the previous reward (r_t−1_) and the previous action (a_t−1_) which is transformed within the hidden layer to output an action in the current trial (a_t_) and an optional state-value estimate (v_t_) (if the loss function is A2C). We systematically trained different network architectures varying in the factors Cell type (RNN or LSTM), Noise (Weber noise or none), Loss (REINFORCE or A2C) and Entropy (none, fixed or annealed) resulting in 24 design combinations (see methods section for details). (b) Example data from a human learner. (c) Example data from an LSTM network with computation noise solving the same task. In b and c, individual choices (colored dots on top) show selected action, and lines denote drifting rewards for each action. % Optimal: Proportion of choices of the most rewarding action. % Switches: Proportion of switches, i.e choice_t_ not equal to choice_t−1_.

### Human data

For comparison with RNN behavior, we re-analyzed human data from a previous study (placebo condition of (Chakroun et al., 2020), n=31 male participants). Participants performed 300 trials of the four-armed restless bandit task as described in the environment section.

### Computational modeling of behavior

Our model space for RNN and human behavior consisted of a total of 21 models (see Table 1). Each model consisted of two components, a *learning rule* (Delta rule or Bayesian learner) describing value updating, and a *choice rule* mapping learned values onto choice probabilities.

**Table 1.**
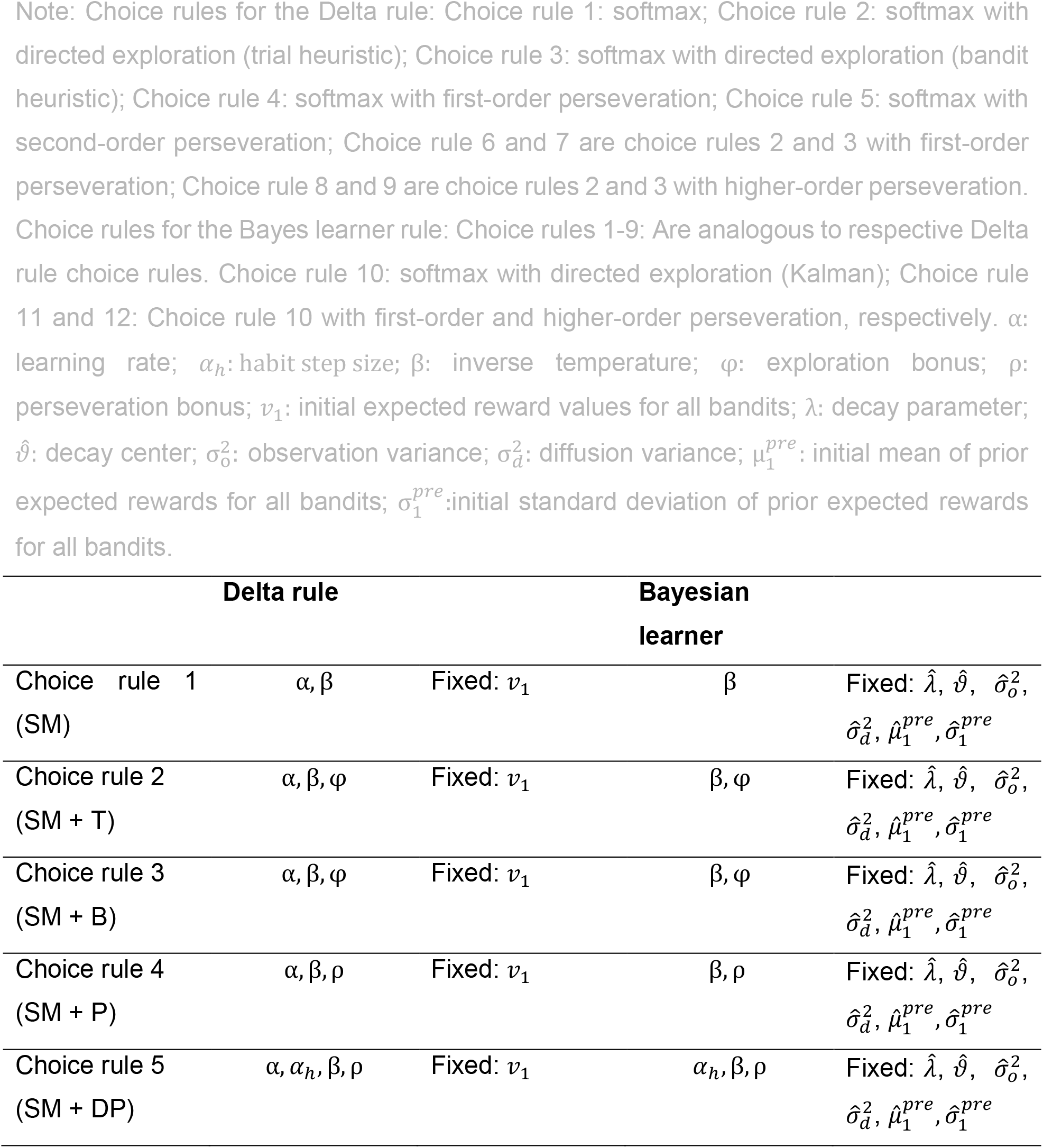
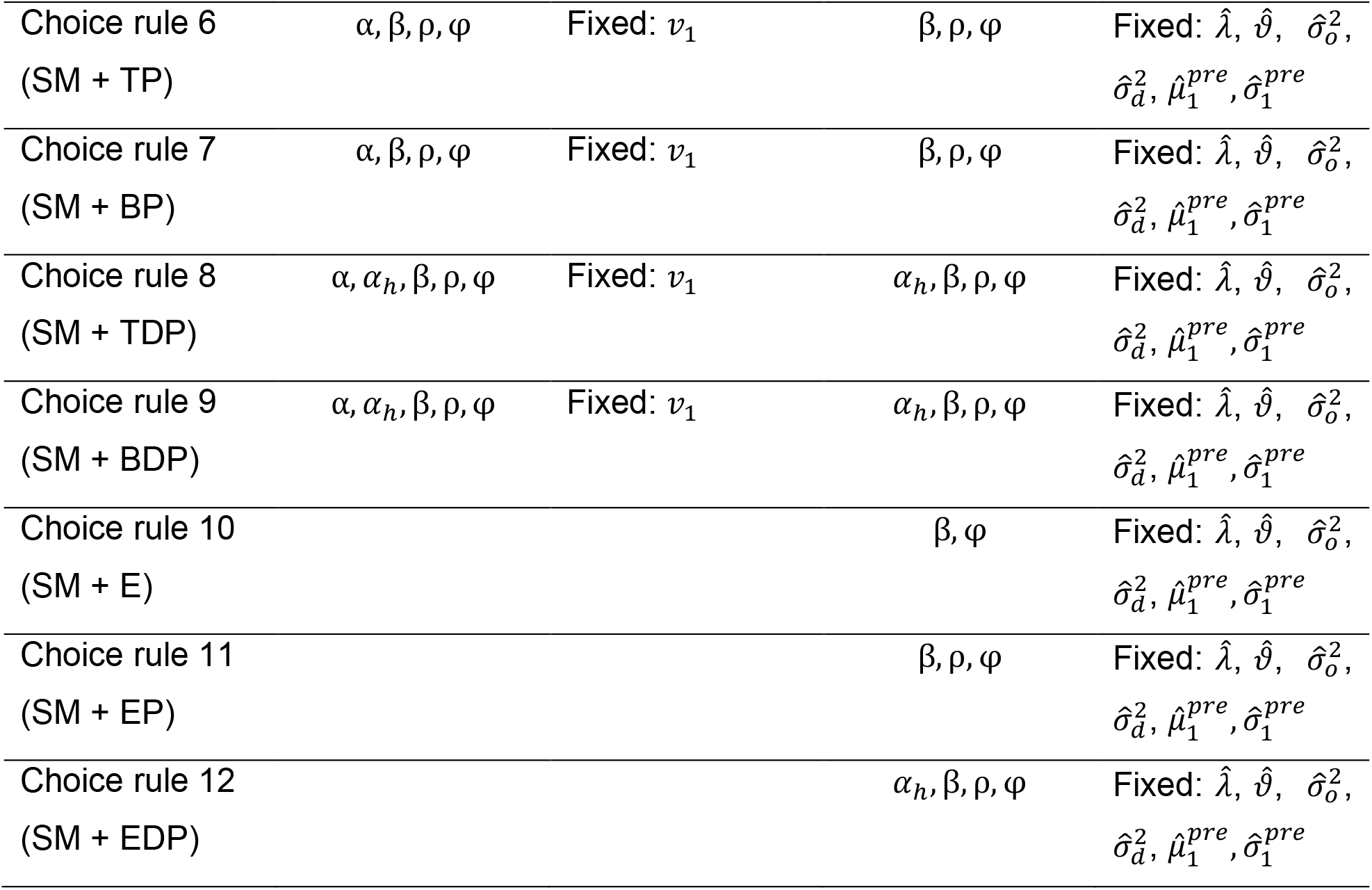
Free and fixed parameters of all computational models.

*Delta rule:* Here, agents update the expected value 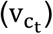 of the bandit chosen on trial t (c_t_) based on the prediction error (*δ*) experienced on trial t:

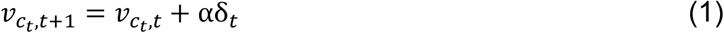

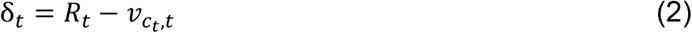

The learning rate 0 ≤ α ≤ 1 controls the fraction of the prediction error used for updating, and *R*_*t*_ corresponds to the reward obtained on trial *t*. Unchosen bandit values are not updated between trials and thus remain *unchanged* until a bandit is chosen again. Bandit values were initialized at *v*_1_ = 50.

*Bayesian learner:* Here we used a standard Kalman filter model (Daw et al., 2006; Kalman, 1960), where the basic assumption is that agents utilize an explicit representation of the process underlying the task’s reward structure. The payoff in trial *t* for bandit *i* follows a decaying Gaussian random walk with mean µ_i,t_ and observation variance 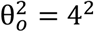. Payoff expectations 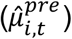 and uncertainties (variances 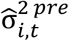) for all bandits are updated between trials according to

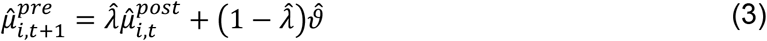

and

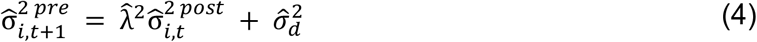

with decay *λ* = 0.9836, decay center *ϑ* = 50 and diffusion variance 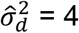.

The chosen bandit’s mean is additionally updated according to

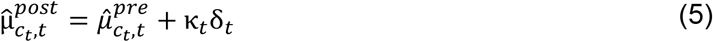

with

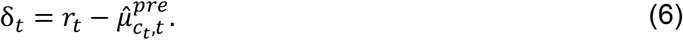

Here, *k* denotes the Kalman gain that is computed for each trial *t* as:

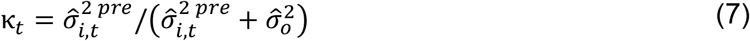

*k*_*t*_ determines the fraction of the prediction error that is used for updating. In contrast to the learning rate in the delta rule model, *κ*_*t*_varies from trial to trial, such that the degree of updating scales with a bandit’s uncertainty 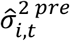. The observation variance 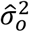 indicates how much rewards vary around the mean, reflecting how reliable each observation is for estimating the true mean. Initial values 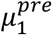 and 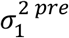 were fixed to 50 and 4 for all bandits, respectively. Estimates of the random walk parameters 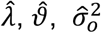 and 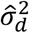 were fixed to their true values (see Table 1).

### Choice rules: Delta rule models

Choice rule 1 used a standard softmax function (SM):

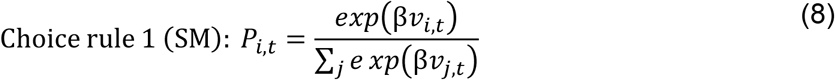

Here, *P*_*i,t*_ denotes the probability of choosing bandit *i* on trial *t* and β denotes the inverse temperature parameter controlling the degree of choice stochasticity.

Choice rule 2 extended choice rule 1 with a heuristic directed exploration term:

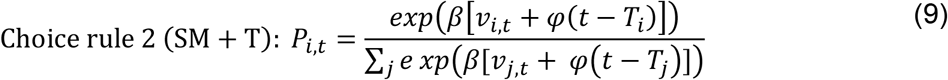

This simple “trial heuristic” (Speekenbrink & Konstantinidis, 2015) models a bandit’s uncertainty as linearly increasing with the number of trials since it was last selected (*t* − *T*_*i*_), where *T*_*i*_ denotes the last trial before the current trial *t* in which bandit *i* was chosen. The free parameter *φ* models the impact of directed exploration on choice probabilities.

Choice rule 3 then replaced the trial-heuristic with a directed exploration term based on a “bandit identity” heuristic:

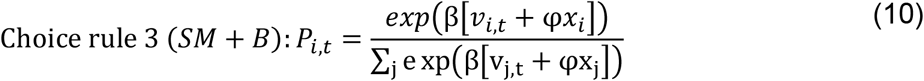

Here, *x*_*i*_ denotes how many unique bandits were sampled since bandit *i* was last sampled. E.g., *x*_*i*_ = 0 if bandit *i* was chosen on the last trial, and *x*_*i*_ = 1 if one other unique bandit was selected since *i* was last sampled. *x*_*i*_ therefore, ranges between 0 and 3.

Choice rule 4 then corresponds to choice rule 1 with an additional first-order perseveration term:

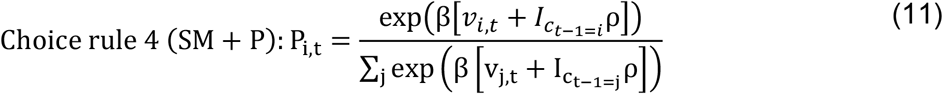

The free parameter ρ models a perseveration bonus for the bandit selected on the preceding trial. *I* is an indicator function that equals 1 for the bandit chosen on trial *t* − 1 and 0 for the remaining bandits.

Choice rule 5 then corresponds to choice rule 1 with an additional higher-order perseveration term:

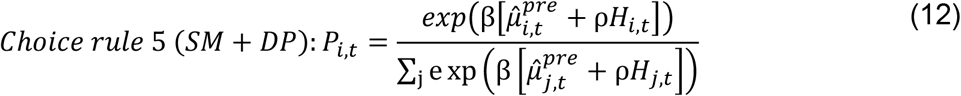

With

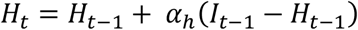

Here, the indicator function *I* is used to calculate the habit strength vector *H*_*t*_ (Miller et al., 2019). *H*_*t*_ is a recency-weighted average of past choices, were the step size parameter *α*_*h*_ [0,1] controls how much recent actions influence current habit strength. In case of *α*_*h*_ = 1, SM+DP corresponds to the first-order perseveration model SM+EP, as only the most recent (previous) action is updated in the habit strength vector. Values of 0 < *α*_*h*_ < 1 result in recency weighted average values for each action in H, giving rise to higher-order perseveration behavior. The resulting habit strength values of each action are then weighted by the perseveration parameter ρ, and enter into the computation of choice probabilities.

Choice rules 6 and 7 likewise extend choice rules 3 and 4 with first-order perseveration terms:

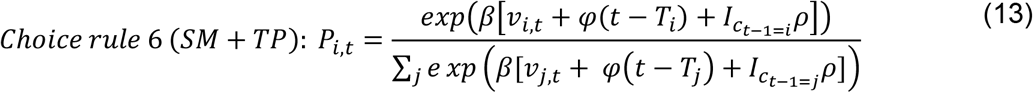

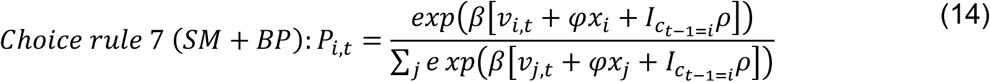

Choice rules 8 and 9 extend choice rules 3 and 4 with higher-order perseveration terms:

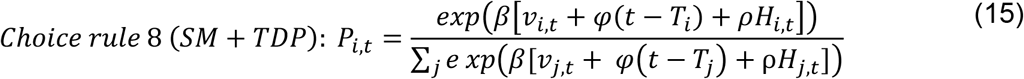

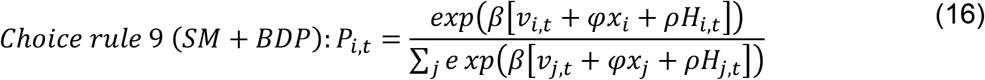

### Choice rules: Bayesian learner models

Substituting*v*_*i,t*_ with 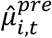 in Equations 8 - 16 yields choice rules 1-9 for the Kalman filter models (equations omitted for brevity). Given that the Bayesian Learner models include an explicit representation of uncertainty, we included two additional models:

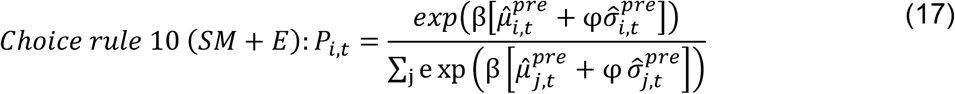

Here, *φ* denotes the exploration bonus parameter reflecting the degree to which choice probabilities are influenced by the uncertainty associated with each bandit, based on the model-based uncertainty 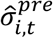. Again, including first order perseveration yields choice rule 11:

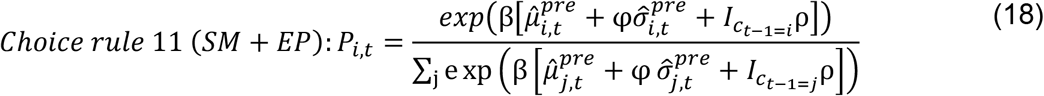

Including higher-order perseveration yields choice rule 12:

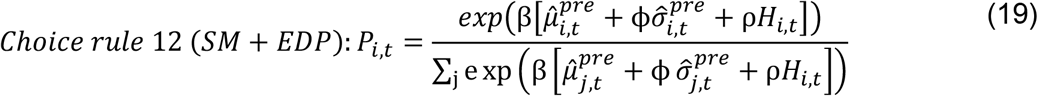

### Model estimation and comparison

Models were fit using Stan and the rSTAN package (Stan Development Team, 2022) in R (Version 4.1.1, R Core Team, 2022). To fit single subject models to human and RNN data, we ran 2 chains with 1000 warm-up samples. Chain convergence was assessed via the Gelman-Rubin convergence diagnostic 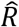 (Gelman & Rubin, 1992) and sampling continued until 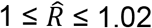 for all parameters. 1000 additional samples were then retained for further analysis.

Model comparison was performed using the loo-package in R (Vehtari et al., 2022) and the Widely-Applicable Information Criterion (WAIC), where lower values reflect a superior model fit (Vehtari et al., 2017). WAICs were computed for each model and human subject/ RNN instance. RNN model comparison focused on the model architecture with the lowest cumulative regret (see Eq. 20). For visualization purposes, we calculated delta WAIC scores for each model by first summing WAIC values for each model over all participants/RNN instances and then subtracting the summed WAIC value of the winning model (Model with the lowest WAIC value if summed over all participants/RNN instances).

### Parameter recovery

We performed parameter recovery for the winning model for human learners (SM+EDP) by simulating a dataset with 100 subjects each performing 300 trials of the 4-armed restless bandit task. The true data generating parameter values were sampled from normal distributions with plausible mean and standard deviation (Danwitz et al., 2022):

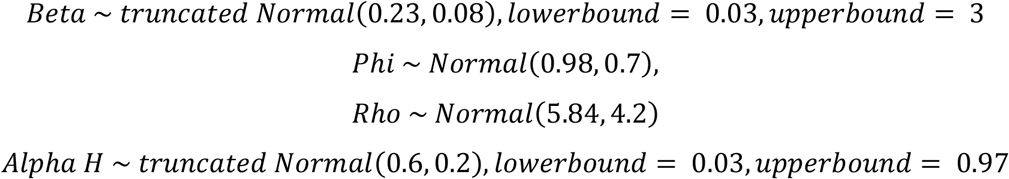

Simulated data were then re-fitted with the best-fitting model (SM+EDP) using the procedures outlined in the model estimation section. The correlation between the true and estimated parameters were taken as a measure of parameter recovery.

#### Cumulative regret

Task performance was quantified using *cumulative regret*, i.e. the cumulative loss due to the selection of suboptimal options, a canonical metric to compare RL algorithms in machine learning (Agrawal & Goyal, 2012; Auer et al., 2002; Wang et al., 2018). Formally, this corresponds to the difference between the reward of the optimal action 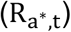 and the obtained reward 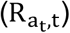, summed across trials:

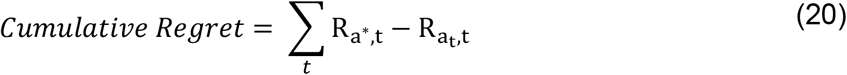

Lower cumulative regret corresponds to better performance. Note that human-level cumulative regret implies that an agent solved the exploration-exploitation tradeoff with human-level performance. Because other measures such as switch rates do not directly reflect performance, we refrained from using such measures as metrics for RNN architecture selection.

#### Code & Data Availability

Code (Python code for the RNN, Stan code for the computational models of behavior) and behavioral data of human and RNN (winning architecture) agents will be shared publicly upon publication.

For review purposes use the following view-only link: https://osf.io/ndpbw/?view_only=a24df99b080f4fe0821a4bff65dc8c47

## Results

### Model-agnostic behavioral results

Our first aim was to identify the best-performing RNN architecture, as deep learning algorithms can be sensitive to hyperparameter settings (Haarnoja et al., 2019; Henderson et al., 2019). The factors considered in the RNN model space are summarized in Table 2 in the Appendix. Performance asymptote was reached by each architecture, such that there was no further improvement from 49.500 to 50.000 training episodes (all *p* > .2)). According to cumulative regret (Fig. 7 in the Appendix), the best-performing architecture used LSTM units in combination with computation noise (Findling & Wyart, 2020), and no entropy regularization during training (Wang et al., 2018). All subsequent analyses therefore are focused on this architecture.

**Table 2.** Overview of factors that are systematically explored in RNN training. Total number of RNN models: 2 (Unit type) x 2 (Computation noise) x 3 (Entropy cost) x 2 (Learning algorithm) = 24 RNN models.

| Factor |  |
| --- | --- |
| Unit type | Vanilla / LSTM |
| Computation noise | None / Update-dependent |
| Entropy cost $\beta_e$ | None / Fixed / Annealed |
| Learning algorithm | REINFORCE / A2C |

**Table 3.** Hyperparameter values used during RNN training.

| Hyperparameters |  |
| --- | --- |
| Number of hidden units | 48 |
| Learning rate | 0.0001 |
| State-value estimate cost $\beta_v$ | 0.5 |
| Weber fraction $\zeta$ | 0.5 |
| Discount factor | 0.5 |
| Training Episodes | 50.000 |
| Trials per Episode | 300 |

We next calculated cumulative regret for each agent (30 RNN instances of the best performing RNN, 31 human subjects from the placebo condition of Chakroun et al., 2020) solving the identical bandit problem (see methods). A Bayesian t-test on the mean cumulative regret on the final trial showed moderate evidence for comparable performance of RNNs and human subjects (*BF*_01_ = 4.274, Fig. 3a). This was confirmed when examining the posterior distribution of the standardized effect size, which was centered at zero (*M* = −0.008, Fig. 3b).

**Fig. 3.**
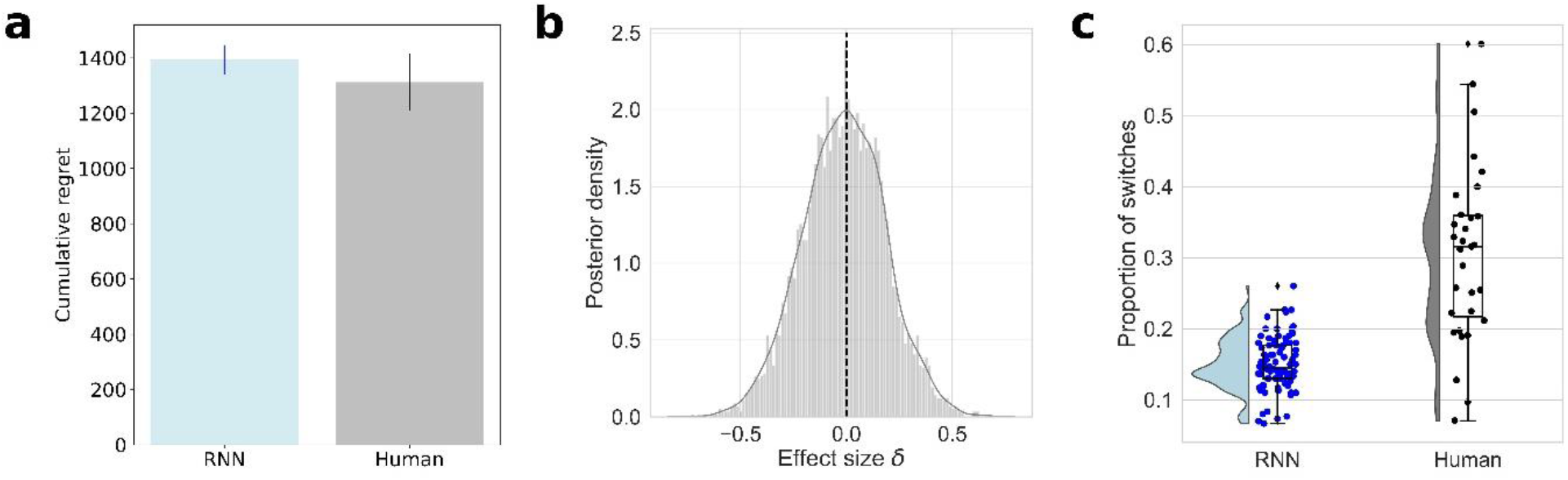
Behavioral data for LSTM networks with computation noise (“RNN”, blue) and human learners ((Chakroun et al., 2020), Placebo condition, black). (a) Mean (+/-SEM) cumulative regret over trials for RNNs (blue) and human learners (black) (b) Posterior distribution of the standardized effect size (δ, Bayesian T-Test) showing moderate evidence against a difference in cumulative regret between RNNs and human learners (BF_01_ = 4.274).(c) Proportion of switches for RNNs (blue) and human learners (black).

Analysis of switching behavior revealed that RNNs switched substantially less than human subjects (Bayesian Mann-Whitney U-Test *BF*_10_ > 100, Median switch probability: 31.5% (human), 14.5% (RNN), Fig. 3c).

### Model comparison

To better understand human and RNN performance on this task, a total of 21 RL models (see methods section) were fitted to the behavioral data. All models were fitted to individual agent data (RNN instances from the best-performing architecture and human data from the placebo condition of (Chakroun et al., 2020)) via Hamiltonian monte Carlo as implemented in STAN. Model comparison was carried out using the Widely Applicable Information Criterion *WAIC* (Vehtari et al., 2017) by computing *ΔWAIC* scores for each model and agent (see methods section), yielding values of 0 for the best-fitting model. The best-fitting model differed for RNN and human agents. Whereas the Kalman-Filter model with higher-order perseveration (SM+DP) was the best model for RNNs, human data were better accounted for by the same model with an additional directed exploration parameter (SM+EDP, see also Table 4 and Table 5 in the Appendix).

**Table 4.**
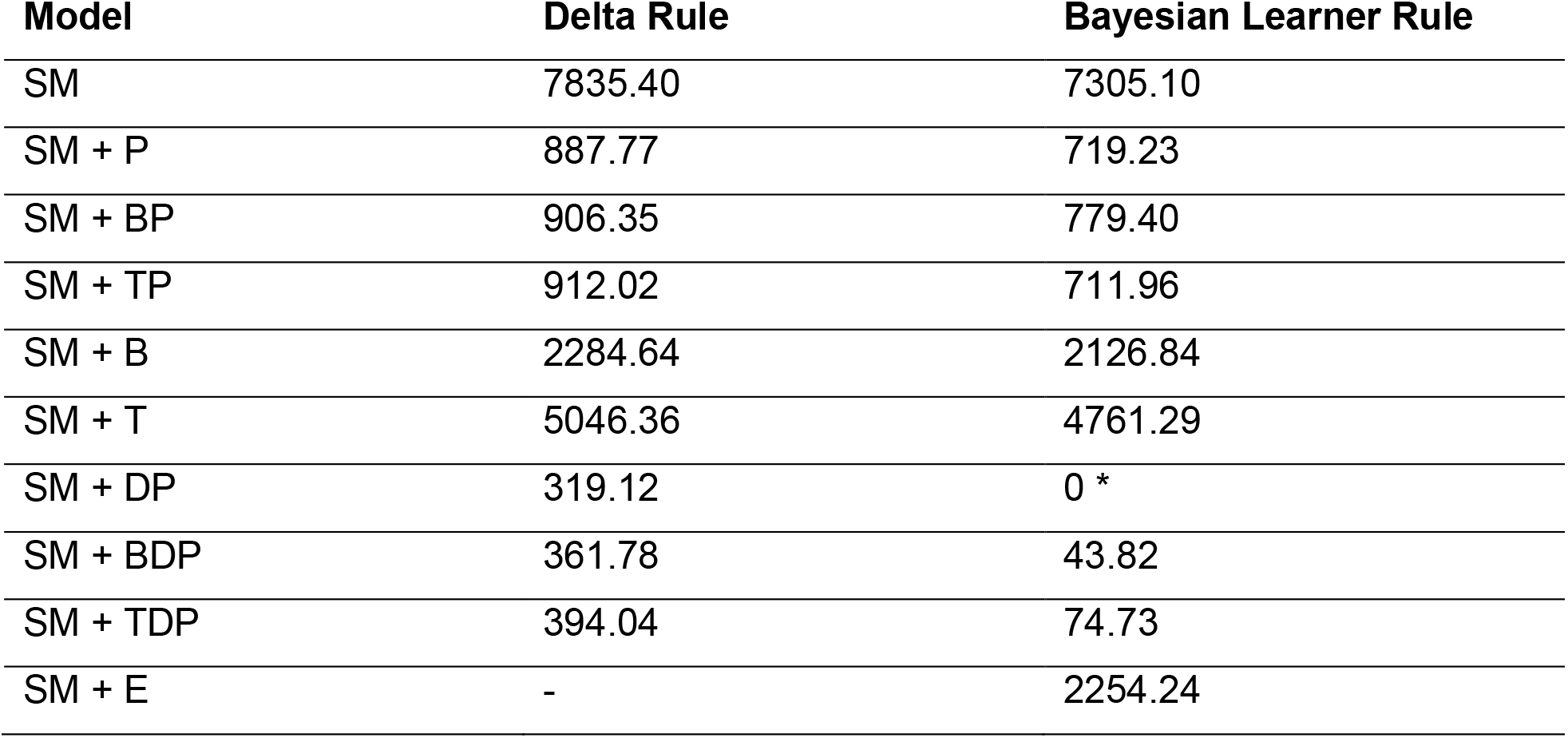
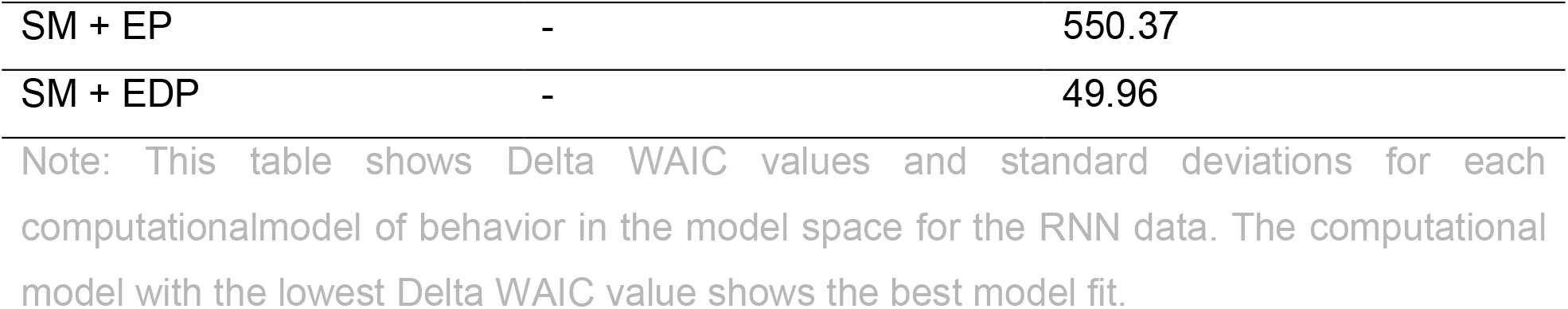
Model Comparison for the best performing RNN architecture (LSTM network with computational noise)

**Table 5.** Model Comparison for human data.

| Model | Delta Rule | Bayesian Learner Rule |
| --- | --- | --- |
| SM | 1057.70 | 967.71 |
| SM + P | 708.45 | 658.87 |
| SM + BP | 388.13 | 333.93 |
| SM + TP | 333.39 | 267.68 |
| SM + B | 858.05 | 767.47 |
| SM + T | 762.96 | 639.96 |
| SM + DP | 646.96 | 575.79 |
| SM + BDP | 243.94 | 230.92 |
| SM + TDP | 236.94 | 196.55 |
| SM + E | - | 655.61 |
| SM + EP | - | 165.43 |
| SM + EDP | - | 0 * |

Note: This table shows Delta WAIC values and standard deviations for each computational model of behavior in the model space for the human data. The computational model with the lowest Delta WAIC value shows the best model fit.

As can be seen from Fig. 4, in RNNs, model fit of the SM+DP (without directed exploration terms) and SM+EDP model (with directed exploration terms) was highly similar. Note, the more basic SM+DP is a nested version of the SM+EDP model with the directed exploration parameter *φ* = 0. In such cases, a straightforward approach is to examine the more complex model, as all information about the nested parameter (here the directed exploration parameter *φ*) is contained in that model’s posterior distribution (Kruschke, 2015). All further analyses therefore focused on the SM+EDP model.

**Fig. 4.**
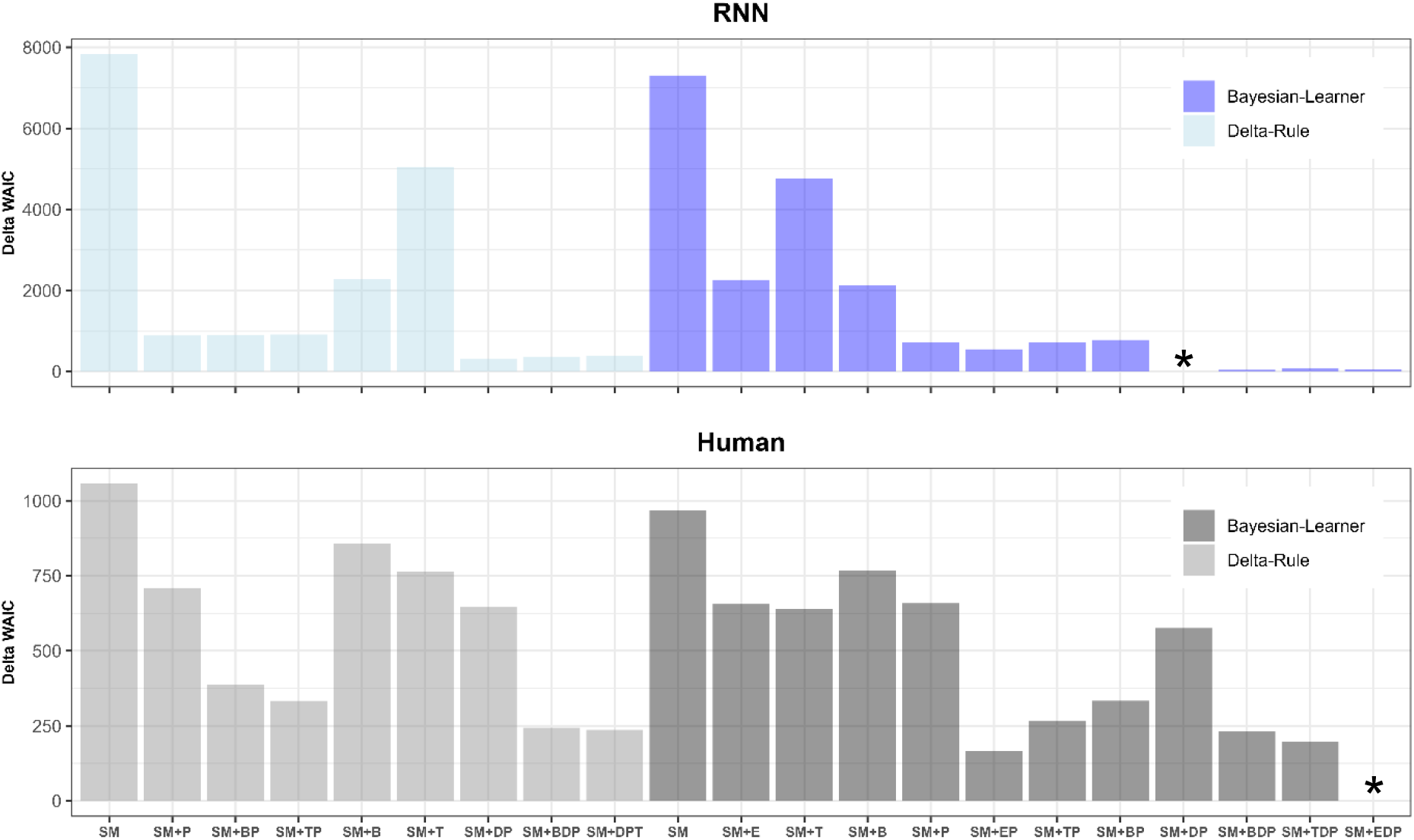
Model comparison via ΔWAIC (see methods section) where smaller values indicate a superior fit. For LSTM networks with computation noise (top panel), the Bayesian learner with higher-order perseveration (SM+DP) accounted for the data best. For human learners (bottom panel), the same model, but including a uncertainty term (SM+EDP) fitted the data best (See star symbols).

To quantify absolute model fit, the posterior predictive accuracy of the SM+EDP model was examined for each agent. Five-hundred data sets were simulated from the model’s posterior distribution, and the proportion of trials in which the simulated and observed choices were consistent were computed and averaged across simulations. Bayesian Mann-Whitney U-Test showed very strong evidence that predictive accuracy (Fig. 5a) was higher for RNNs than for human learners (RNN: *Mdn* = 0.793, *range*: [0.675,0.895], Human: *Mdn* = 0.666, *range*: [0.385,0.904], *BF*_10_ > 100).

**Fig. 5.**
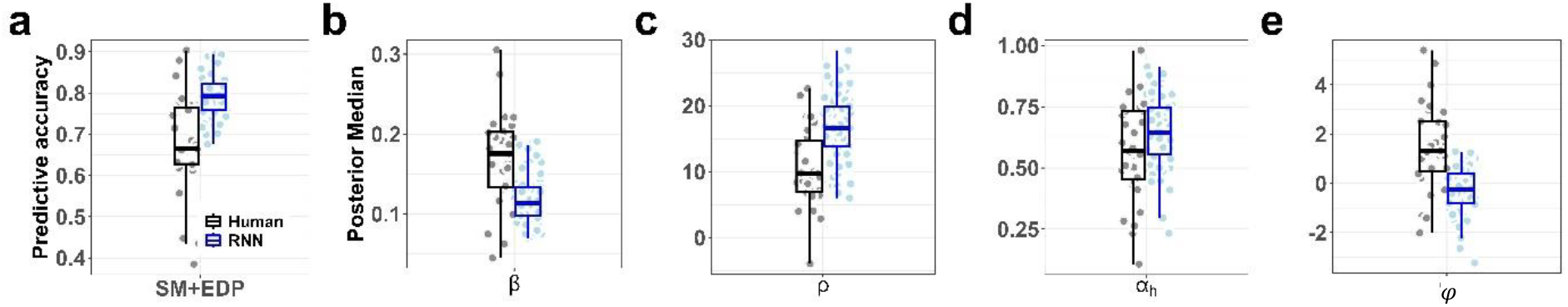
Median posterior values of model parameters for the best-fitting model (Bayesian learner with exploration and higher order perseveration terms, SM+EDP). (a) Predictive accuracy as the fraction of choices correctly predicted by the fitted computational model of behavior, (b) choice stochasticity parameter β, (c) perseveration parameter ρ, (d) Habit step size parameter α_h_, (e) exploration parameter φ for human learners (black) and RNNs (blue).

**Fig. 6.**
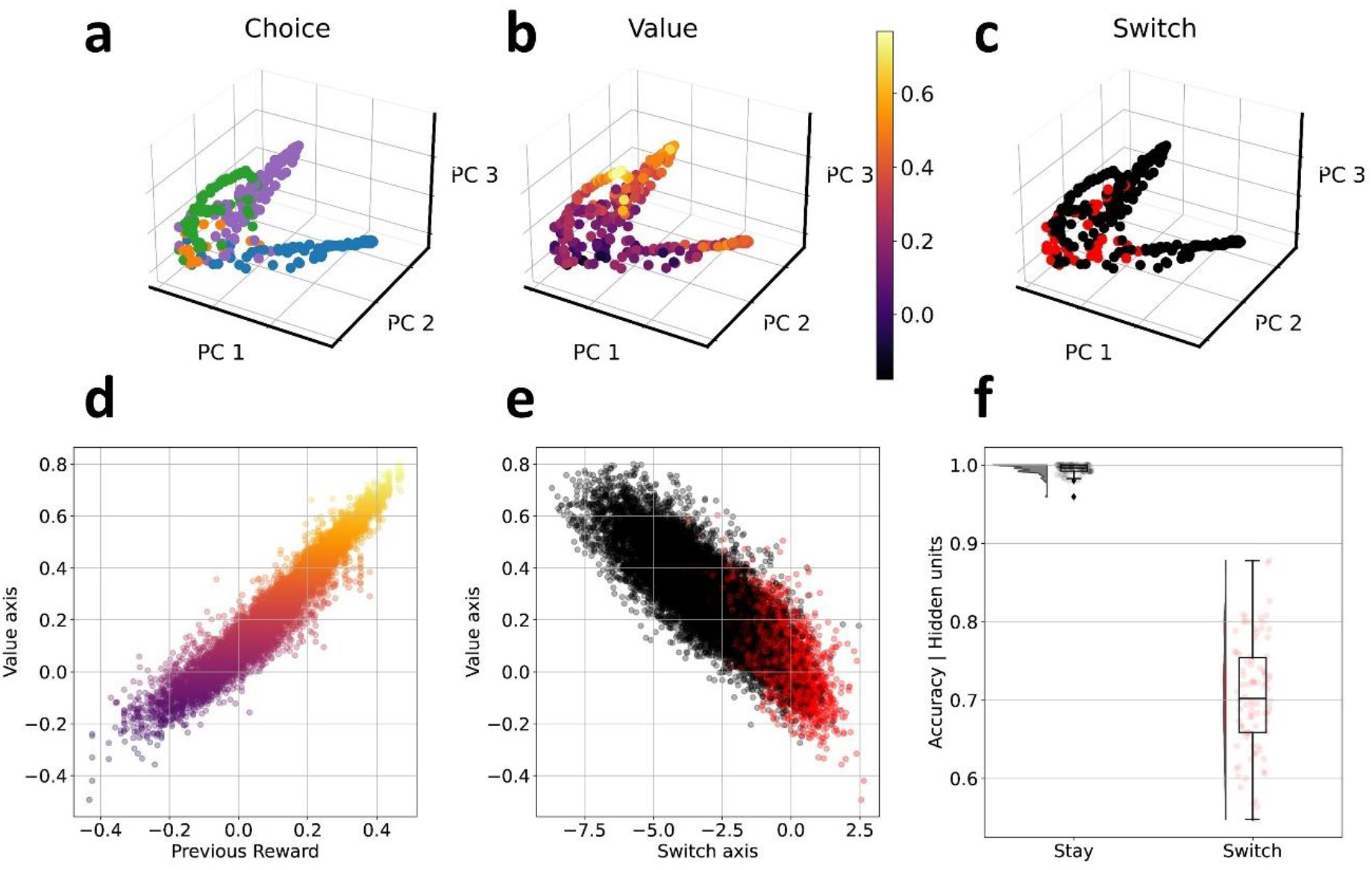
Hidden unit activation dynamics for a single network instance (a-c) and across all instances (d, e). **a-c:** Hidden unit dynamics (first three principal components) of an example RNN agent color coded by choice (a), state-value estimate (b) and switching behavior (c, switch – red, stay – black). (d, e): Targeted dimensionaility reduction. (d) The state-value axis (y-axis) was highly correlated with previous reward (x-axis). Note that previous reward is a continuous variable in the range -0.5 and 0.5 (as rewards in the range: [0,1] were mean centered). (e) Lower state-value (y-axis) was linked to greater log-odds of switching (x-axis). (f) Accuracy of choice prediction given the PCA-based de-noised hidden unit activation state using a multinomial model revealed almost perfect accuracy for stay decisions (99%) and reduced, but above chance-level accuracy for switch decisions (70%).

### Parameter recovery

We ran parameter recovery simulations to ensure that our modeling approach could correctly identify the data generating process (Wilson & Collins, 2019) (see Parameter recovery section in the methods and Fig. 8 in the Appendix). This revealed good parameter recovery for most parameters of the SM+EDP model (all correlations between true and estimated parameter values are r > .76), with the exception of α_h_ (habit step size, r = .492). Therefore, some caution is warranted when interpreting this parameter.

**Fig. 7.**
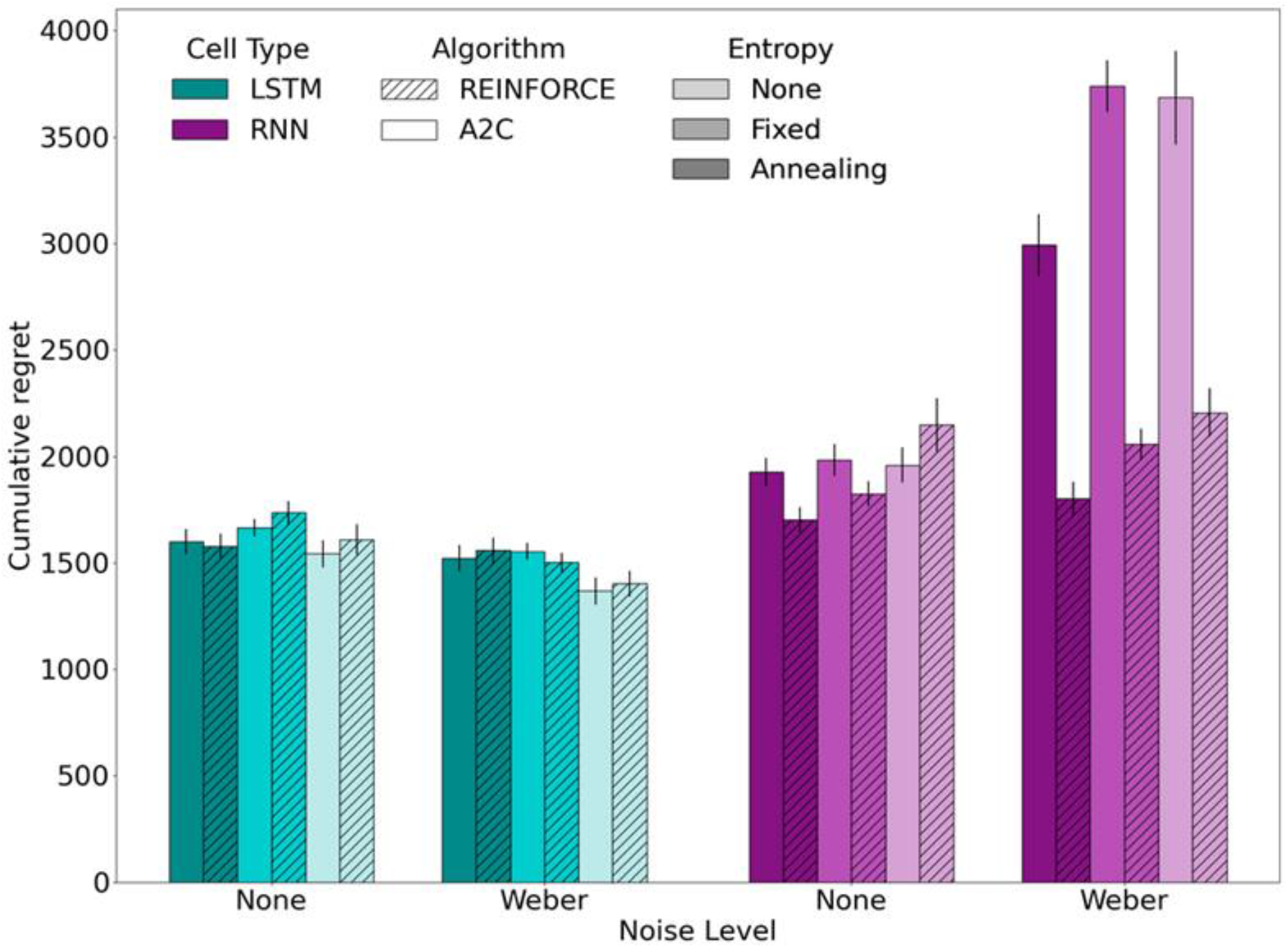
Mean final cumulative regret of all RNN architectures in the restless bandit task by different design factors. X-axis denote whether no computation noise (“None”) or weber noise (“Weber”) is added to hidden units. Error bars denote SEM.

**Fig. 8.**
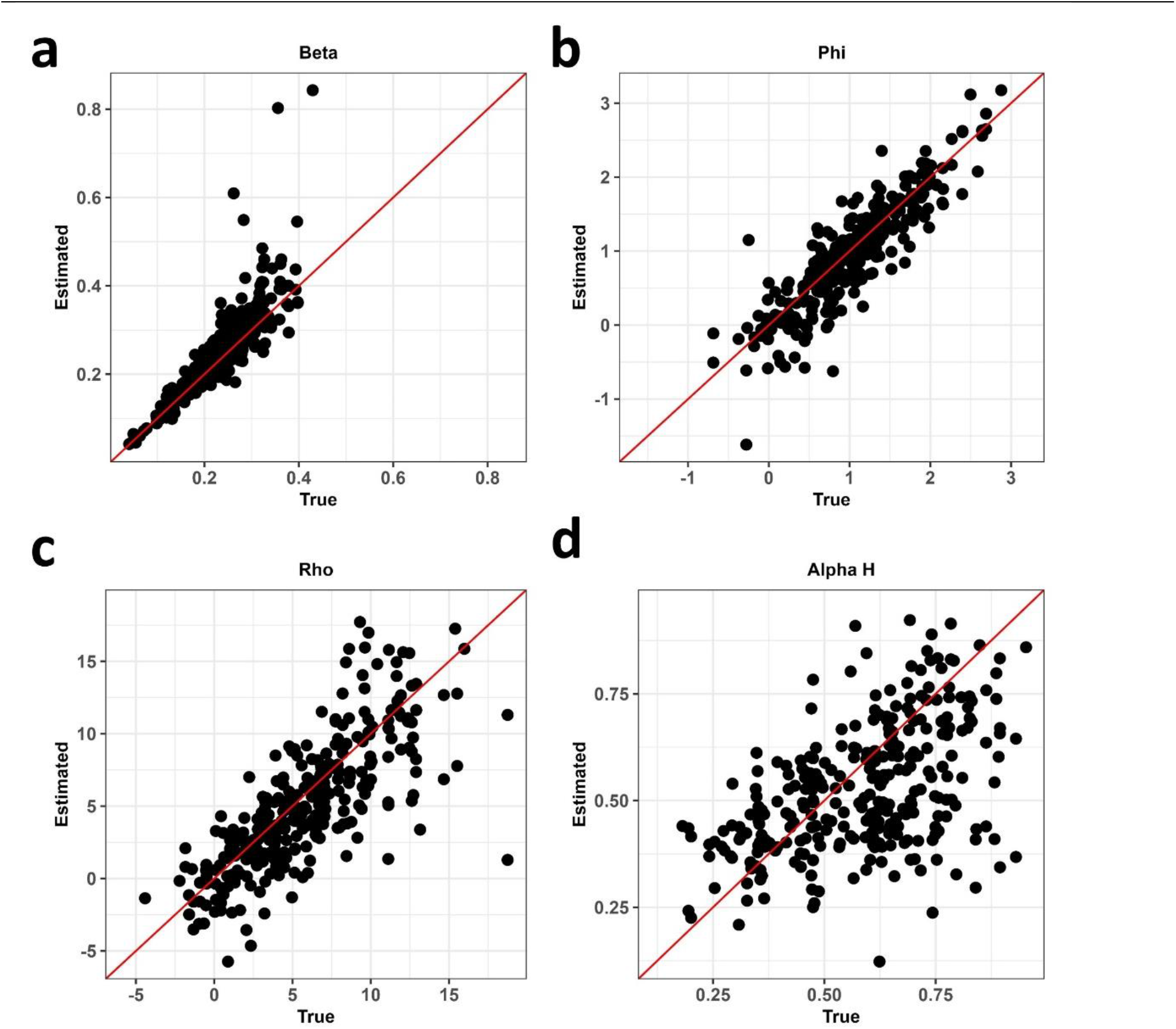
Parameter recovery. (a) Beta (choice stochasticity), (b) Phi (exploration bonus), (c) Rho (perseveration bonus), (d) Alpha H (habit step size). The scatterplots show the correlation of the true data generating parameter values and estimated values after fitting the model to the generated data. Dots correspond to the median of the posterior distribution.

### Posterior predictive checks

Posterior predictive checks were conducted by simulating five-hundred data sets from the best-fitting model for each human learner and RNN agent. While the model reproduced the qualitative differences between human learners and RNNs (simulated switch proportions were higher for humans compared to RNNs) the model somewhat overpredicted switch proportions in both cases (See Fig. 9 in the Appendix).

**Fig. 9.**
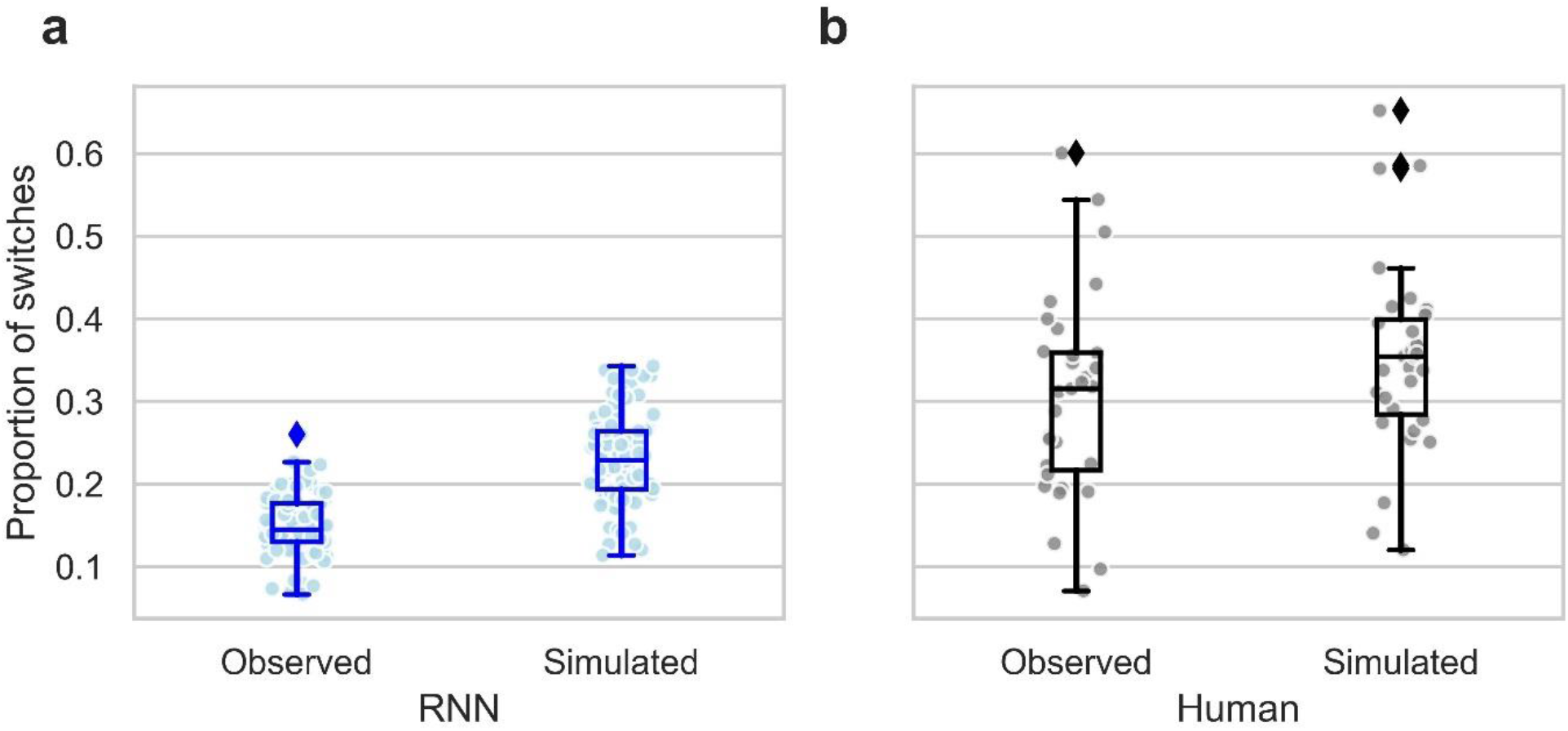
Posterior predictive checks: Comparison of observed and simulated switch proportions from the SM+EDP model. A: RNN agents, B: Human learners. Simulations reproduce qualitative differences between RNN agents and human learners, but overpredict switch proportions for both, quantitatively. Five-hundred data sets for each human learner and RNN agent were simulated from the model’s posterior distribution, and the proportion of switch trials were computed. Simulated data points refer to the median switch rate across the five-hundred data sets resulting in one value for each observed switch rate.

### Analysis of model parameters

Taking this into account, we examined the model parameters (medians of individual subject posterior distributions) of the best-fitting model and compared them between human learners and RNNs, again focusing on the SM+EDP model (for corresponding results for the SM+DP model, see Fig. 10 in the Appendix). All subsequent Bayes Factors (*BF*) are based on the Bayesian Mann-Whitney U-Test (van Doorn et al., 2020) testing for differences in median values between RNNs and human learners. Choice consistency (β) was lower for RNNs than human learners (Fig. 5b, RNN: *Mdn* = 0.113, *range*: [0.0698, 0.191], Human: *Mdn* = 0.175, *range*: [0.0448, 0.305], *BF*_10_ = 91.3). RNNs showed substantially higher levels of perseveration (Fig. 5c, RNN: *Mdn* = 16.6, *range*: [6, 28.4], Human: *Mdn* = 9.66, *range*: [−3.97, 22.6], *BF*_10_ = 76.23), but there was only anecdotal evidence for a greater habit step size than human learners (Fig. 5d, RNN: *Mdn* = 0.644, *range*: [0.233, 0.913], Human: *Mdn* = 0.568, *range* [0.108, 0.980], *BF*_10_ = 2.31). In line with the model comparison (see above) there was very strong evidence for reduced directed exploration in RNNs compared to human learners (Fig. 5e, RNN: *Mdn* = −0.234, *range*: [−3.23, 1.29]), Human: *Mdn* = 1.32, *range*: [−2.01, 5.40], *BF*_10_ > 100). Bayesian Wilcoxon Signed-Rank Tests (van Doorn et al., 2020) nonetheless showed strong evidence for *φ* estimates < 0 in RNNs (*BF*_10_ > 19.98) whereas *φ* estimates were reliably > 0 in human learners (*BF*_10_ > 100).

**Fig. 10.**
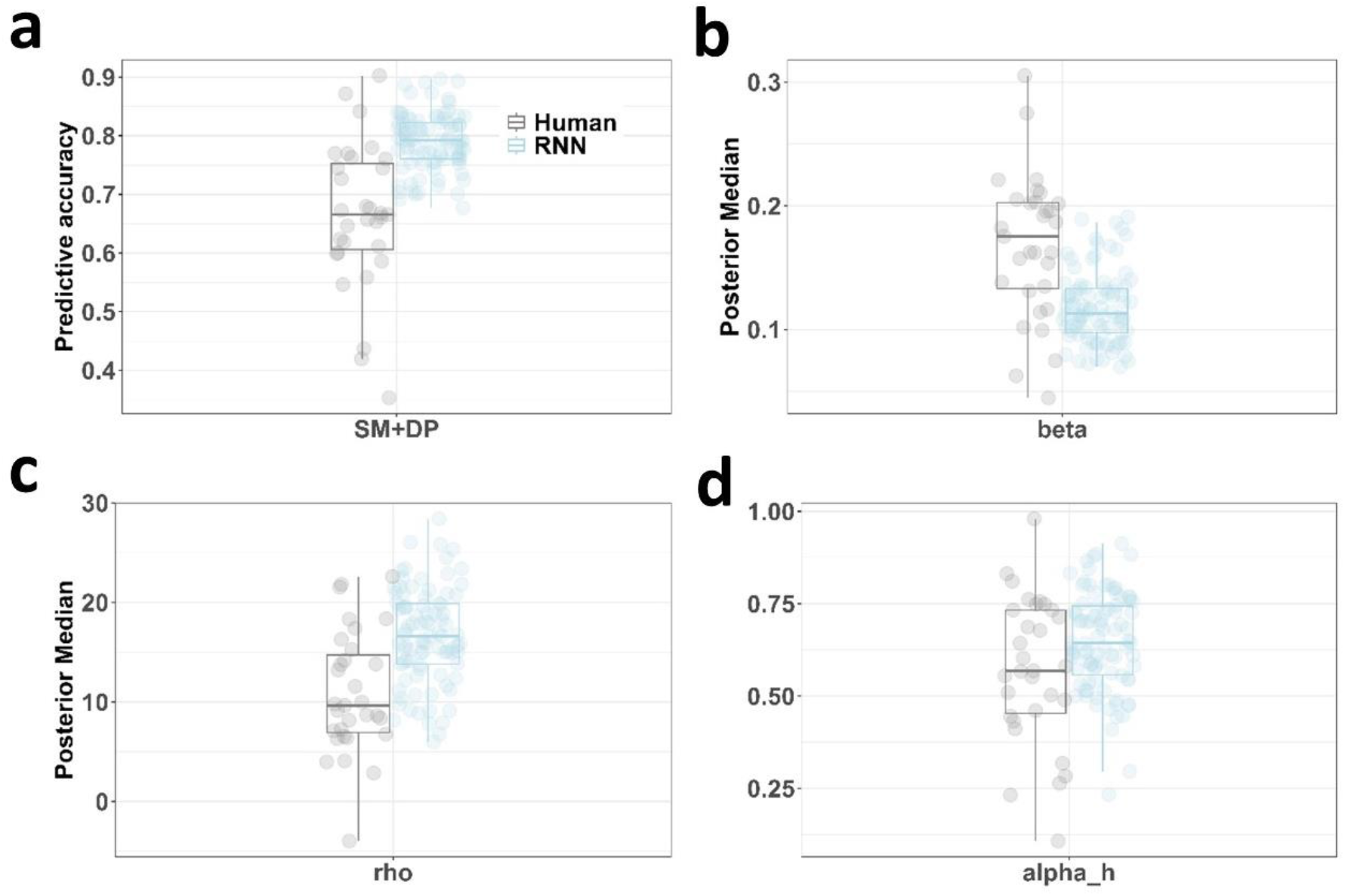
Median posterior values of model parameters for the best-fitting model for RNN agents (Bayesian learner with higher order perseveration term, SM+DP). (a) Predictive accuracy as the fraction of choices correctly predicted by the computational model of behavior, (b) choice stochasticity parameter β, (c) perseveration parameter ρ, (d) Habit step size parameter α_h_.

Taken together, both the model comparison and the analysis of parameter estimates show that human learners, but not RNNs, adopted a directed exploration strategy. In contrast, RNNs showed substantially increased perseveration behavior.

### Hidden unit analysis

Finally, we investigated RNN hidden unit activity. This analysis is similar to the analysis of high dimensional neural data (Cunningham & Yu, 2014) and we first used PCA to visualize network dynamics. The first three principal components accounted for on average 73% of variance (see Fig. 11 in the Appendix). The resulting network activation trajectories through principal component space were then examined with respect to behavioral variables. Coding network state by behavioral *choice* revealed separated choice-specific clusters in principal component space (see Fig. 6a for hidden unit data from one RNN instance, and see Fig. 12 in the Appendix for the corresponding visualizations for all instances). The degree of spatial overlap of the choice-specific clusters appeared to be related to the state-value estimate of the RNN (Fig. 6b), with greater overlap for lower state values. Coding network state according to stay vs. switch behavior (repeat previous choice vs. switch to another bandit, as a raw metric for exploration behavior, Fig. 6c) revealed that switches predominantly occurred in the region of activation space with maximum overlap in choice-specific clusters, corresponding to low state-value estimates. Highly similar patterns were observed for all RNN instances investigated (see Fig. 12 in the Appendix).

**Fig. 11.**
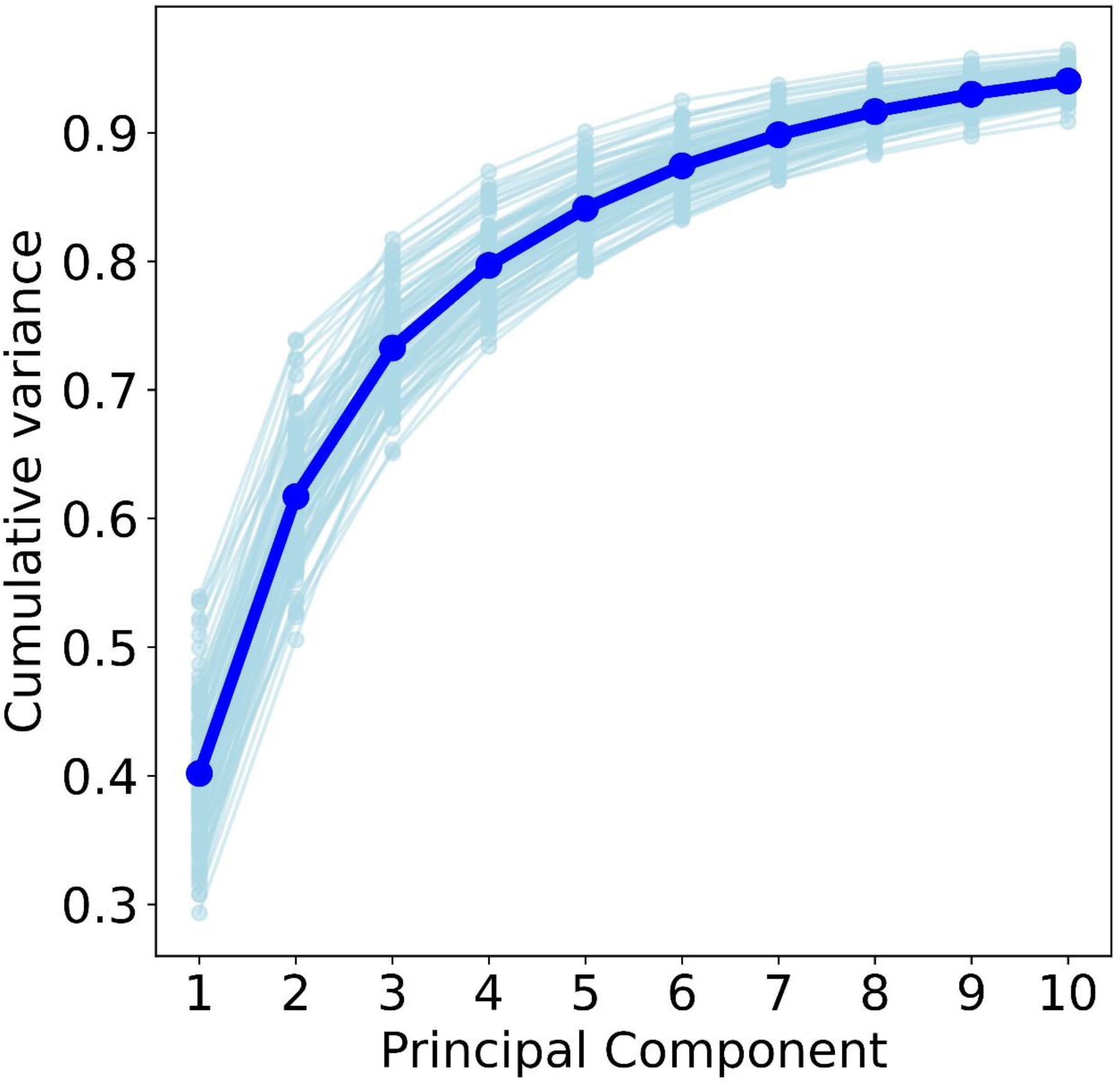
Cumulative variance explained in hidden unit activity by principal components of the best RNN architecture (LSTM cell with weber noise, A2C algorithm and no entropy regularization): Light blue lines denote cumulative variance explained for each RNN instance. Dark blue line denotes mean cumulative variance explained over all RNN instances.

**Fig. 12.**
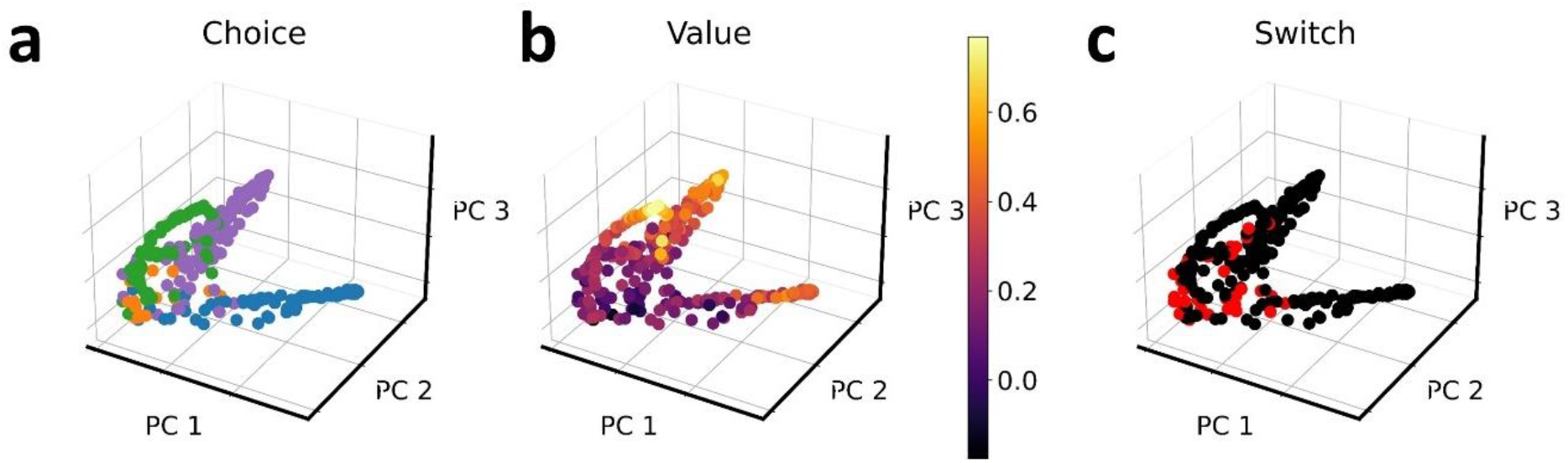
Hidden unit dynamics (first three principal components) of an example RNN agent color coded by choice (a), state-value estimate (b) and switching behavior (c, switch – red, stay - black). The pattern of choice-specific clusters (a), more overlap for actions corresponding to low state-value estimates (b) and switches occuring predominantly in this region of overlap (c) generalizes to all RNN instances under consideration (see below)

One downside of the PCA-analysis is that components are not readily interpretable, and, due to differential rotation of these patterns in PC-space, is not straightforward to conduct analyses across all network instances. We address these issues via targeted dimensionality reduction (TDR) (Mante et al., 2013), which projects the PCA-based hidden unit data onto novel axes with specific interpretations (see methods section). We first used TDR to project the PCA-based hidden unit data onto a *value axis*, i.e., the predicted state-value estimate given the hidden unit activity on a given trial. Across all network instances, predicted state value was highly correlated with the reward obtained by the network on the previous trial (*r*(17998) = .97), Fig. 6d). We next explicitly tested the visual intuition that switching behavior predominantly occurs when state value is low, and projected the hidden unit data onto a *switch axis* via logistic regression (see methods section), corresponding to the log-odds of observing a switch. Positive log-odds indicated that a switch decision is more likely than a stay decision, and vice versa for negative log-odds. Results confirmed the results from the analysis of single network instances (e.g., Fig. 6b, 6c): switches predominantly occurred when estimated state value was low, as reflected in a negative correlation of the *value axis* and the *switch axis* scores (*r*(17998) = −.80, Fig. 6e).

Further, we asked whether switching occurs randomly, or follows a predictable pattern. To this end, a multinomial model was fitted to predict RNN choices given the current PCA-based de-noised hidden unit activity. We then compared the accuracy of choice prediction between stay and switch trials. If RNNs follow a switching strategy, the accuracy of predicting switching decisions from de-noised hidden unit activity should be above chance level (0.25). The prediction accuracy was near perfect for stay decisions (*M* = 0.996, see Fig. 6f) and markedly disrupted but still substantially above chance level for switch decisions (*M* = 0.702, see Fig. 6f). This is consistent with the idea that RNNs rely on more than choice randomization to solve the exploration-exploitation dilemma.

## Discussion

Here we comprehensively investigated exploration mechanisms in recurrent neural network (RNN) models trained to solve restless bandit problems, reinforcement learning tasks commonly used in cognitive and systems neuroscience. We expanded upon previous work in four ways. First, in contrast to earlier work (Findling & Wyart, 2020; Wang et al., 2018) we focused on four-armed restless bandit problems, allowing for a more comprehensive analysis of exploration behavior. Second, we systematically investigated a range of RNN design choices and resulting impacts on performance. Third, we directly compared human and RNN behavior, both in terms of performance (cumulative regret) and using computational modeling, when solving the exact same task problem. Finally, we investigated exploration mechanisms in the best-performing network architecture via a comprehensive analysis of hidden unit activation dynamics.

We extensively tested and transparently report upon a total of twenty-four RNN design factor combinations. The architecture exhibiting the best performance was an LSTM network, combined with computation noise as previously suggested (Findling & Wyart, 2020), no entropy regularization and trained with the advantage-actor-critic (A2C) algorithm (Mnih et al., 2016). The superior performance of the LSTM versus the vanilla RNN is not surprising. LSTMs are endowed with a more sophisticated memory process (Equations 25 – 29) where different gating mechanisms regulate the impact of past experiences (previous actions and rewards) on current choices. These mechanisms allow LSTM networks to learn dependencies over longer time scales than vanilla RNNs (Hochreiter & Schmidhuber, 1997). The performance benefit of LSTMs also resonates with a well-known computational model of learning and action selection mechanisms in prefrontal cortex (PFC) and basal ganglia (O’Reilly & Frank, 2006). This model is characterized by a combination of LSTM-like gating mechanisms that are combined with an Actor-Critic architecture. Here, the *Actor* (i.e., the basal ganglia) updates actions by gating working memory updating processes in the PFC. The *Critic* (i.e., midbrain DA neurons) estimates reward values of possible actions, thereby adjusting future actions to maximize reward. Similar to Findling & Wyart (Findling & Wyart, 2020) our results show the benefit of biologically-inspired computation noise (“Weber Noise”, (Findling & Wyart, 2020)) added to hidden unit activity that scales with the degree of recurrent activity reconfiguration between subsequent trials. Entropy regularization, in contrast, adds noise to the policy during training to discourage premature convergence to a suboptimal policy (Mnih et al., 2016). This “update-dependent” noise mechanism might thus entail exploration-specific noise that contrasts with the introduction of general stochasticity as implemented in entropy regularization schemes. This result further reinforces the performance-enhancing effect of computation noise observed in human reward-guided decision making (Findling et al., 2019) which is thought to be modulated by reciprocal connections of the locus coeruleus-norepinephrine system and the anterior cingulate cortex (Findling & Wyart, 2021; McClure et al., 2005). This also resonates with results from deep neural networks, where various noise-based schemes have been implemented to improve network performance and/or to increase the resilience of the networks under sparse information, e.g. noise at the input level, similar to dataset augmentation methods (Goodfellow et al., 2016), at the level of the weight update (An, 1996) or during computation (Dong et al., 2020; Fortunato et al., 2019; Qin & Vucinic, 2018).To clarify differences between human and RNN behavior, we applied comprehensive computational modeling of behavior (Farrell & Lewandowsky, 2018). Model comparison according to WAIC revealed that human behaviour was best accounted for by a Bayesian learning model (Kalman filter) with an uncertainty-based directed exploration term and higher-order perseveration (SM+EDP). This model is an extension of a previously applied model (SM+EP) (Beharelle et al., 2015; Chakroun et al., 2020; Wiehler et al., 2021) with higher-order perseveration according to a formalism suggested by Miller et al. (Miller et al., 2019), and similar to Lau & Glimcher 2005. RNN behaviour, on the other hand, was best accounted for by the same model without a directed exploration term, although the numerical differences in WAIC to models with directed exploration were numerically very small.

In such situations where model comparison yields somewhat inconclusive results for a full model (SM+EDP) and nested versions of this model (e.g. SM+DP corresponds to SM+EDP with *φ* = 0), there are different possible ways to proceed. The first would be to only examine the best-fitting model (SM+DP). Alternatively, one could examine the full model (SM+EDP), as it contains the nested model as a special case, and allows to directly quantify the degree of evidence that e.g. *φ* = 0 (Kruschke, 2015). After all, the posterior distribution of *φ* contains all information regarding the value of this parameter, given the data and the priors. Following suggestions by Kruschke (Kruschke, 2015), we therefore focused on the full model. Indeed, this analysis revealed strong evidence that *φ* < 0 in RNNs, highlighting the caveats of solely relying on categorical model comparison in cases of nested models.

Both RNNs and human learners exhibited higher-order perseveration behavior, i.e., perseveration beyond the last trial. This tendency was substantially more pronounced in RNNs than in human learners, and is conceptually linked to exploration, because perseveration can be conceived of as an uncertainty-avoiding strategy. However, similar to analysis using first-order perseveration (Chakroun et al., 2020), accounting for higher-order perseveration did not abolish the robust evidence for directed exploration in our human data. This is in line with the idea that human learners may apply perseveration (uncertainty avoiding) and directed exploration (information gain) strategies in parallel (Payzan-LeNestour et al., 2013). RNNs showed substantially higher levels of perseveration than human learners. Perseveration is often thought to be maladaptive, as learners “stick” to choices regardless of reward or task demands (Dehais et al., 2019; Hauser, 1999; Hotz & Helm-Estabrooks, 1995) and it is a hallmark of depression and substance use disorders (Zuhlsdorff, 2022), behavioral addictions (de Ruiter et al., 2009), obsessive compulsive disorder (Apergis-Schoute & Ip, 2020) and tightly linked to PFC functioning (Goldberg & Bilder, 1987; Munakata et al., 2003). In the light of these findings, it might appear surprising that RNNs show a pronounced tendency to perseverate. But perseveration might support reward accumulation by enabling the network to minimize losses due to excessive exploration (see below). Another take-away from this study could be that RNNs and human learners use perseveration to trade-off between maximizing reward and minimizing policy complexity as was discussed in previous research (Gershman, 2020).

Analysis of model parameters also revealed strong evidence for uncertainty aversion in RNNs (even when accounting for higher-order perseveration), reflected in an overall *negative* exploration bonus parameter *φ* (i.e., an “exploration malus”). In contrast, human learners showed the expected positive effect of uncertainty on choice probabilities (directed exploration) (Chakroun et al., 2020; Schulz et al., 2019; Schulz & Gershman, 2019; Wiehler et al., 2021; Wilson et al., 2014, 2021). Importantly, this divergence between human and RNN mechanisms demonstrates that directed exploration is not required for human-level task performance. The observation of information seeking behavior (directed exploration) independent of reward maximization can be understood as “non-instrumental” information-seeking, which was shown in many human studies (Bennett et al., 2021; Bode et al., 2023; Brydevall et al., 2018). Our results further validate that for human learners, in contrast to RNN agents, information is intrinsically rewarding, independent of the accumulation of external reward(Bennett et al., 2016).Overall, human learners perseverate less and explore more (Fig. 3c), thereby potentially avoiding the costs of prolonged perseveration and finding the optimal bandit faster by continuously exploring the environment. Both strategies converge on comparable performance. Finally, RNNs showed a lower inverse temperature parameter (β) than human learners. All things being equal, lower values of β reflect more random action selection, such that choices depend less on the terms included in the model. A β-value of zero would indicate random choices, and as β increases, the policy approaches a deterministic policy in which choices depend completely upon the model terms. However, the absolute level of choice stochasticity reflected in a given value of β also depends on the absolute values of all terms included in the model. Whereas the absolute magnitude of value and exploration terms was comparable between human and RNNs, the perseveration term was about twice the magnitude in RNNs, which explains the lower β-values in RNNs. The results from the analysis of predictive accuracy also confirmed that a greater proportion of choices was accounted for by the best-fitting computational model in RNNs compared to humans, showing that these differences in β do not reflect a poorer model fit.

The computational models used here have conceptual links to two other models popular in RL and computational neuroscience, upper confidence bound models (UCB, (Auer et al., 2002)) and Thompson sampling (Thompson, 1933). In UCB, uncertainty affects action value estimates additively (Schulz & Gershman, 2019), similar to the uncertainty bonus implementations used in in this as well as previous work (Chakroun et al., 2020; Daw et al., 2006; Speekenbrink & Konstantinidis, 2015; Wiehler et al., 2021). In UCB, the uncertainty bonus is a function of the number of trials since an action was last selected (similar to the trial heuristic in Eq. 9) and the current trial number. In Thompson sampling, uncertainty affects action value estimates multiplicatively (Schulz & Gershman, 2019), similar to the inverse temperature parameter in the softmax function. Actions have probability distributions (e.g., gaussian with mean corresponding to the current action value, and variance corresponding to action value uncertainty), which are updated between subsequent trials. The Kalman Filter models in this work also implement such Bayesian updating. In pure Thompson sampling action values are sampled from these probability distributions with more or less width according to uncertainty, which results in more or less stochastic action selection. Our Kalman Filter implementations do not use such sampling, but select actions based on current mean (action value estimates, 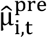) and add current variance 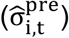 as an uncertainty bonus more similar to UCB.

To investigate the computational dynamics underlying RNN behaviour, we initially applied dimensionality reduction of hidden unit activations patterns via Principal Component Analysis (PCA) (Findling & Wyart, 2020; Mante et al., 2013; Wang et al., 2018). The first three principal components accounted for on average 73% of variance in hidden unit activity (see Fig. 11 in the Appendix). Visual inspection of activation patterns in principal component space then revealed three effects: First, coding network state by behavioral *choice* revealed clearly separated choice-specific clusters in principal component space, an effect that was observed across all RNN instances examined (see Fig. 12 in the Appendix). Second, the degree of spatial overlap of the choice-specific clusters directly related to the state-value estimate of the network. Action representations on trials with higher state-value estimates were more separated than during trials with lower state-value estimates. Again, this pattern was observed across all RNN instances examined (see Fig. 12 in the Appendix) and resonates with systems neuroscience work showing neural populations are more predictive for high-value actions than for low-value actions (Ebitz et al., 2018). Oculomotor regions like the frontal eye field (FEF) (Ding & Hikosaka, 2006; Glaser et al., 2016; Roesch & Olson, 2003, 2007) and the lateral intraparietal area (LIP) (Platt & Glimcher, 1999; Sugrue et al., 2004) show more pronounced choice-predictive activation patterns during saccades to high vs. low value targets. Third, to investigate the link between RNN dynamics and exploration, we coded network state in PC-space by stay vs. switch behavior. This revealed that switches predominantly occurred in the region of activation space with maximum overlap in choice predictive clusters, corresponding to low state-value estimates. Again, this effect was observed across all network instances examined. Generally, these observations show that 1) switches occurred predominantly during trials with low state value, 2) low state value was associated with less pronounced choice-predictive activation patterns. Although these patterns were qualitatively highly similar across RNN instances (see Fig. 12 in the Appendix), the geometrical embedding of these effects in principal component space differed. This illustrates one downside of PCA the components as such are not directly interpretable, and the different rotations of patterns in principal component space complicate the aggregation of analyses across network instances.

To address this issue, and to obtain interpretable axes, we applied targeted dimensionality reduction (TDR) (Ebitz et al., 2018; Mante et al., 2013). TDR projects the PCA-based de-noised hidden unit activation patterns onto novel axes with clear interpretations (see methods section), allowing for a quantification of the intuitions gained from PCA. We projected the de-noised hidden unit data onto a *value axis*, i.e., the predicted state-value estimate given the de-noised hidden unit activity on a given trial. Across all network instances, this measure was highly correlated with the reward obtained on the previous trial (Fig. 6d). Likewise, we projected the de-noised hidden unit data onto a *switch-axis*, i.e., the predicted log-odds of observing a switch, given the de-noised hidden unit activity on a given trial. Across all network instances, this axis showed a strong negative correlation with the value-axis, confirming that indeed the log-odds of switching increased with decreasing state value, and decreased with increasing state value, resembling a Win-Stay-Lose-Shift (WSLS) strategy (Herrnstein, 1997) that accounts for substantial choice proportions also in human work (Worthy et al., 2013). However, pure WSLS would predict much higher switch rates than observed in RNNs, suggesting that RNNs show a mixture of a WSLS-like strategy in conjunction with high perseveration.

Finally, we decoded choices from de-noised hidden unit activation dynamics, and compared prediction accuracy for stay vs. switch decisions. The decoder showed near perfect accuracy for stay decisions, which resonates with animal work showing that neural choice decoding is improved during perseveration (Coe et al., 2002; Ebitz et al., 2018). Importantly, performance of the decoder was lower, but still substantially above chance-level for switches. The de-noised hidden units therefore represent an activation pattern that can be utilized to correctly predict switch-targets, suggesting that switching behavior is not entirely based on choice randomization.

One caveat of this work is that, although often applied in the context of RL in volatile environments (Domenech et al., 2020; Kovach et al., 2012; Swanson et al., 2020), the comparison between stay and switch trials does not unequivocally map onto the exploitation vs. exploration distinction. For example, stay decisions can be due to greedy choices (choosing the option with the highest expected reward) but also due to perseveration. In contrast, switch decisions can be due to random or strategic exploration (Wilson et al., 2021) and may involve more complex model-based strategies and/or simpler heuristics like following motor patterns such as exploring by choosing each available option once and then exploiting (Fintz et al., 2022). We nonetheless applied the stay vs. switch distinction, as it makes by far the least assumptions regarding what constitutes exploration vs. exploration.

Several limitations of this work need to be addressed. First, our conclusions are restricted to the specific type of RNN architectures and task family studied here. Other network architectures may use different computational mechanisms to solve exploration-exploitation problems. Although our final network model space resulted in a total of 24 different RNN architectures, the impact of additional design choices such as network size, learning rate, discount factor, type of activation function or values for the Weber fraction (noise) were not systematically explored. Although the combination of LSTM with the A2C algorithm is robust to different hyperparameter settings (Mnih et al., 2016), a different RNN architecture or hyperparameter combination could have yielded even better performance or could have produced a form of directed exploration. Future research could benefit from the use of other architectures such as transformer models (L. Chen et al., 2021; Parisotto et al., 2020; Upadhyay et al., 2019) or explore the role of these additional factors. Second, a general limitation of this approach more generally is that neural network models, although roughly based on neuronal computations, suffer from a number of biological implausibilities (Pulvermüller et al., 2021). These include the backpropagation algorithm used to update the parameters of the network (Lillicrap et al., 2020), the lack of separate modules analogous to different brain regions, and lack of neuromodulation mechanisms (Pulvermüller et al., 2021). However, some recent work has begun to address these shortcomings (Mei et al., 2022; Robertazzi et al., 2022). Third, it can be argued that the restless bandit task is a suboptimal paradigm to investigate directed exploration, because reward and uncertainty are confounded, unlike other tasks, such as the horizon task (Wilson et al., 2014). But the episodic character of this task (a series of discrete games where reward-related information is reset for every episode) has the caveat that explore-exploit behaviour can only unfold over a relatively small time frame (1-6 trials), whereas in the restless bandit task it can evolve over much longer time periods (e.g. 300 trials). Fourth, we studied RNNs in a meta-RL framework, by training them to solve tasks from a given task family, and investigating the mechanisms underlying their performance. Other work has used RNNs in a supervised manner to predict human behavior (RNNs essentially replaced the computational models) (Dezfouli et al., 2019; Eckstein et al., 2023; Fintz et al., 2022; Ger et al., 2024). Future studies might compare RNNs as models for human behavior to the computational models examined here. Further, the behavioral signatures we observed in human learners and RNN agents could additionally or alternatively be affected by value decay, i.e. forgetting or decay of Q-values due to limited cognitive resources, in particular in human learners (Collins & Frank, 2012; Niv et al., 2015). Note that such a value decay mechanism is explicitly included in the Kalman Filter model (i.e. via the decay center ϑ and decay rate λ). However, we held these parameters fixed to their true values for all models, as estimating them is notoriously difficult due arising convergence issues (Wiehler et al., 2021). For example, even in hierarchical models, which are more robust than the individual-participant models that we focus on here, ϑ and λ could only be estimated when they were implemented in a non-hierarchical manner, and not jointly. For this reason, we refrained from examining this issue further. Another caveat is that parameter recovery for the habit step size parameter α_h_ in the SM+EDP model was low (r = .492), such that this parameter must be interpreted with caution, although this habit-strength process was used in computational modeling of behavior in previous work (Gershman, 2020; Lau & Glimcher, 2005; Miller et al., 2019; Palminteri, 2023). In contrast, recovery of the other parameters was excellent, in particular rho and phi (Danwitz et al., 2022), which showed robust differences between RNNs and human learners. Further, in terms of posterior predictive checks the SM+EDP model accurately reproduced the substantial difference in switch rates between RNNs (Fig. 9a in Appendix) and human learners (Fig. 9b in Appendix). In both cases, however, the model overpredicted switch rates. We think that this hints towards that there is still variance left in accounting for stay decisions even if one accounts for choice stochasticity, higher order perseveration and directed exploration. Last, an important limitation is that human behavioral data analyzed here are from german male participants (age 18-35)(Chakroun et al., 2020). Exploration behavior may differ according to gender (C. S. Chen et al., 2021; van den Bos et al., 2013) and age (Mizell et al., 2024; Nussenbaum & Hartley, 2019; Sojitra et al., 2018), and future research would benefit from a more representative sample.

Taken together, we identified a novel RNN architecture (LSTM with computation noise) that solved restless four-armed bandit tasks with human-level accuracy. Computational modeling of behavior showed both human learners and RNNs exhibit higher-order perseveration behavior on this task, which was substantially more pronounced for RNNs. Human learners, on the other hand, but not RNNs, exhibited a directed (uncertainty-based) exploration. Analyses of the networks’ exploration behavior confirmed that exploratory choices in RNNs were primarily driven by rewards and choice history. Hidden-unit dynamics revealed that exploration behavior in RNNs was driven by a disruption of choice predictive signals during states of low estimated state value, reminiscent of computational mechanisms in monkey PFC. Overall, our results highlight how computational mechanisms in RNNs can at the same time converge with and diverge from findings in human and systems neuroscience.

## Declarations Section

### Author Contribution

Conceptualization: DT, IP, JP. Methods: DT, AB, IP, JP. Implementation: DT, IP. Writing (first draft): DT. Writing (review and editing): AB, IP, JP. Supervision: JP. Funding acquisition: JP.

### Ethics Approval and Consent to Participate

Not applicable

## Appendix RNN unit types

The present study systematically compared network architectures consisting of four different types of RNN units, standard recurrent units (“vanilla” units) (Elman, 1990), standard LSTM units (Hochreiter & Schmidhuber, 1997), as well as both unit types endowed with computation noise.

### Vanilla RNN

This is a simple Elman recurrent neural network (Elman, 1990) (“Vanilla” RNN). Let *X*_*t*_ denote the input to the network at time *t, H*_*t*_ the recurrent state of the network and *Y*_*t*_ the output of the network. Then the network is governed by the following equations:

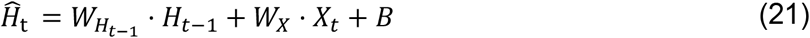

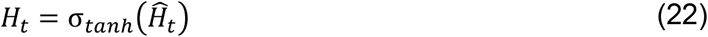

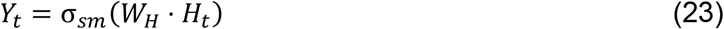

The input *X*_*t*_ is a vector of length 5 with the first element corresponding to the reward observed on the previous trial (*r*_*t*−1_) and the remaining elements containing a one-hot encoding of the previously chosen action (*a*_*t*−1_). The latter is a vector of length 4 (No. of actions) with all elements being “0” except the element which corresponds to the previous action being set to “1”. The parameters of the network are weight matrices 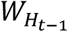, *W*_*X*_ and *W*_*H*_ and the bias vector *B*, which are optimized during the training process. Non-linear activation functions σ_*tanh*_ and σ_*sm*_ denote the hyperbolic tangent and the softmax function, respectively.

The forward pass of the model starts by passing information from the input layer to the hidden layer by calculating the updated state 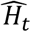 as a linear combination of the input *X*_*t*_ and the previous recurrent activity *H*_*t*−1_ weighed by corresponding weight matrices *W*_*X*_ and 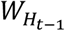 and an additive bias term *B* (Eq. 21). Within the hidden layer, 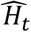 is non-linearly transformed by the hyperbolic tangent function which results in the current recurrent activity *H*_*t*_ (Eq. 22). Recurrent activity *H*_*t*_ in the hidden layer is then transformed to action probabilities *Y*_*t*_ by applying the softmax function to *H*_*t*_ weighed by matrix *W*_*H*_ (Eq. 23).

### Noisy Vanilla RNN

Computation noise might aid in adverse conditions during decision-making (Findling & Wyart, 2020). Following Findling et al. (Findling & Wyart, 2020), we therefore modified the standard RNN unit by adding update-dependent computation noise to the recurrent activity *H*_*t*_.

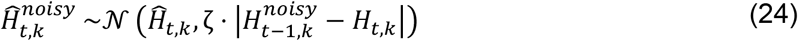

To transform the exact recurrent activity *H*_*t,k*_ in unit *k* in trial *t* to noisy recurrent activity 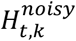 we added noise according to a Gaussian distribution with mean equal to the exact updated state 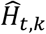 (see Eq. 21) and a standard deviation of the absolute magnitude of the difference between the previous noisy recurrent activity 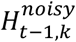 and the current exact recurrent activity *H*_*t,k*_ (see Eq. 22). Because of this difference term the spread of the noise added to a unit scales with the amount of reconfiguration in recurrent activity between subsequent trials similar to a prediction error. The standard deviation is further scaled by the hyperparameter ζ > 0 denoted the *Weber fraction* (Findling & Wyart, 2020). Finally, after having sampled the noisy updated state 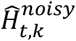 the hyperbolic tangent activation function is applied (see Eq. 22) to calculate noisy recurrent activity 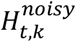. Importantly, similar to Findling et al. (Findling & Wyart, 2020) we treated computation noise as an endogenous constraint to the network where the source of the noise is not modifiable, thereby ignoring the gradients resulting from it in the optimization procedure during gradient descent.

### LSTM

One issue with standard vanilla RNN units is that during gradient descent, they can suffer from exploding and vanishing gradient problems, resulting in the network not learning the task (Rehmer & Kroll, 2020). Long short-term memory networks (LSTMs) can mitigate this problem by using gated units that control the information flow, which allows the network to learn more long term dependencies within the data (Hochreiter & Schmidhuber, 1997).

LSTM behavior is governed by the following standard equations:

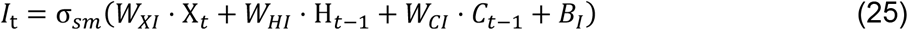

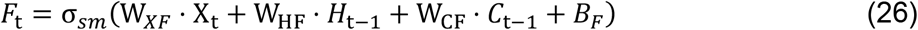

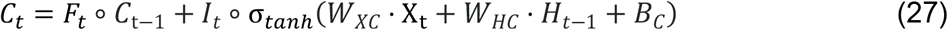

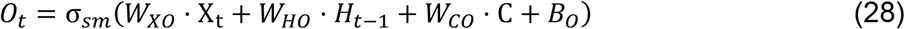

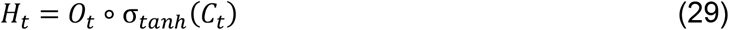

Here, *X* is the same input as in Vanilla RNNs, *I* is the input gate, *F* is the forget gate (also called the maintenance gate), *C* is the cell state, *O* is the output gate, *H* is the hidden state, *t* indexes trials and σ_*sm*_ and σ_*tanh*_ denote the softmax or the hyperbolic tangent activation function, respectively. The trainable parameters of the network are weight matrices *W* and the bias vectors *B*, where subscripts indicate the connected gates/states.

### Noisy LSTM

Following Findling & Wyart (Findling & Wyart, 2020), we introduce Weber noise at the level of the hidden state *H* of an LSTM unit *k* at trial *t*:

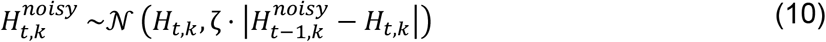

That is, noisy LSTM units are the direct analogue to the noise extension outlined above for vanilla units.

### Training and test environments

Networks were trained and tested on four-armed restless bandit tasks (Daw et al., 2006). For a schematic depiction of the training and testing procedure see Fig. 1. On each trial, agents choose between one of four actions (bandits). Associated rewards slowly drift according to independent gaussian random walks. A single episode during training and testing consisted of 300 trials.

During training, the reward associated with the *i*th bandit on trial *t* was the reward on trial t-1, plus noise:

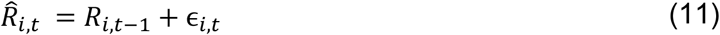

Noise ϵ was drawn from a gaussian distribution with mean 0 and standard deviation 0.1:

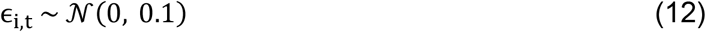

To ensure a reward range: [0,1], reflecting boundaries were applied such that

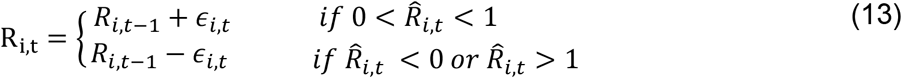

Note that R_i,t_ is a continuous reward as implemented in past research with similar tasks (Chakroun et al., 2020; Daw et al., 2006; Wiehler et al., 2021).

Following training, networks weights were fixed, and performance was examined on three random walk instances previously applied in human work (Chakroun et al., 2020; Daw et al., 2006; Wiehler et al., 2021). Here, the reward associated with the *i*_*th*_ bandit on trial *t* was drawn from a gaussian distribution with standard deviation 4 and mean µ_*i,t*_ and rounded to the nearest integer.

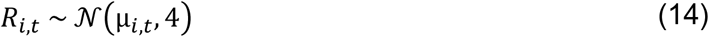

On each trial, the means diffused according to a decaying gaussian random walk:

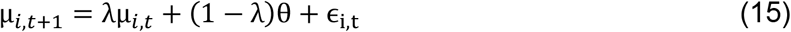

with decay λ = 0.9836 and decay center θ = 50. The diffusion noise ϵ_i,t_ was sampled from a gaussian distribution with mean 0 and SD 2.8:

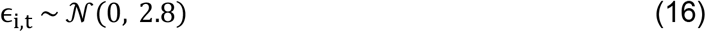

During test, each network was exposed to the same three instantiations of this process used in human work (Chakroun et al., 2020; Daw et al., 2006; Wiehler et al., 2021). Rewards in these three random walks were in the range: [0,100]. As RNNs were trained with rewards in the range: [0,1] we scaled rewards accordingly before testing the RNNs.

### Training procedure

The networks were trained to optimize their weights and biases by completing 50.000 task episodes. For each episode, a new instantiation of the environment was created according to the equations above. We compared two training schemes, the standard REINFORCE algorithm (Williams & Peng, 1991) and advantage actor-critic (A2C, (Mnih et al., 2016)). The objective of the network was to maximize the expected sum of discounted rewards according to following equation:

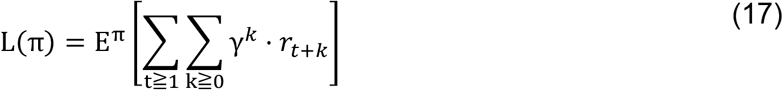

Here, *t* is the trial number, γ the discount factor, *t*_*t*+*k*_ the observed reward at trial *t* + *k* (k is a positive integer) and π is the policy followed by the network.

Following each episode, one of the following algorithms was used to update the network parameters to improve the policy. REINFORCE (Williams & Peng, 1991) relies on a direct differentiation of the objective function:

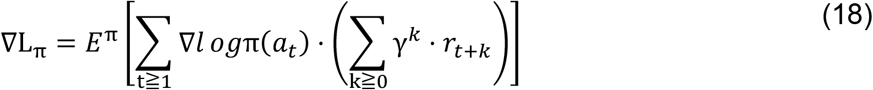

Here the gradient of the policy loss (∇L_π_) is calculated by first generating a single rollout from the current policy (e.g., a trajectory of actions and rewards in an episode), and then summing the derivatives of the log probabilities of chosen actions (Logπ(a_t_)) weighted by the discounted sum of expected rewards from the current trial until the end of the episode. Note that this ensures that action probabilities will increase or decrease according to the expected rewards following these actions. Based on a single rollout from the current policy, if the expected returns (∑_k≧0_ γ^k^ · r_t+k_) are positive, the gradient will be positive and therefore gradient descent will increase the log probabilities of chosen actions. Conversely, if expected returns based on a single rollout from the current policy are negative, the gradient will be negative and therefore gradient descent will decrease the log probabilities of chosen actions. Thus, action probabilities for actions that led to rewards will be increased, and action probabilities for actions that led to punishments will be decreased.

Advantage Actor-Critic (A2C, (Mnih et al., 2016)) uses a weighted sum of the policy gradient (*∇L*_*π*_), the gradient with respect to the state-value function loss (*∇L*_*v*_), and an optional entropy regularization term (*∇L*_*ent*_), defined as follows:

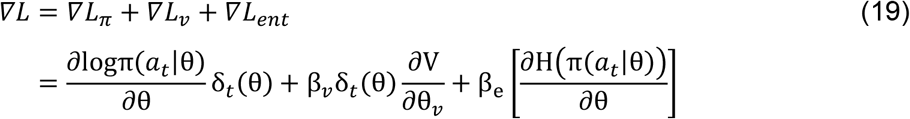

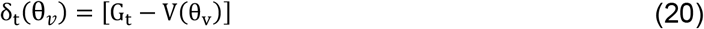

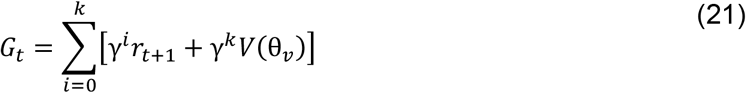

Here, on a given trial *t*, the chosen action is denoted by *a*_*t*_, the discounted return is *G*_*t*_ with *k* being the number of steps until the end of the episode, the actor component with the action policy π is parameterized by RNN parameters θ, the critic component with the value function *V* estimating the expected return is parameterized by θ_*v*_, the entropy of policy π is denoted by *H*(π), the advantage function that estimates the temporal-difference error is denoted by *δ*_*t*_(θ_*v*_). A2C In this formulation contains two hyperparameters β_*v*_ and β_e_ scaling the relative influence of the state-value function loss and the entropy regularization term, respectively.

Note that in equations 39 – 41, RNN parameters corresponding to the policy θ and to the value function θ_*v*_ are separated, but in practice, as in Wang et al. (Wang et al., 2018), they share all layers except the output layer where the policy corresponds to a softmax output and the value function to a single linear output. Weights and biases for REINFORCE and A2C were updated using the RMSProp algorithm as implemented in TensorFlow 1.15.0. We initialized all weights with the standard Xavier initializer (Glorot & Bengio, 2010) as implemented in TensorFlow. Bias parameters were initialized with 0 vectors.

A common problem of policy gradient methods such as REINFORCE is high variance in the gradients used during the stochastic gradient descent (SDG) optimization procedure (Sutton & Barto, 2018). This is the case because the magnitude of the gradients depends on the empirical returns (sum of collected rewards in a given episode). We therefore mean-centered rewards to reduce the variance in the gradients, which improved the training process and performance.

A subset of hyperparameters were systematically varied, as outlined in Table 2. The entropy cost (β_*e*_) was either set to 0.05 (fixed entropy), linearly annealed over the course of training from 1 to 0, or omitted (*none*). In networks with computation noise, the Weber fraction ζ was set to 0.5 (Findling & Wyart, 2020). Additional hyperparameters (learning rate, discount factor, no. of training episodes etc.), were selected based on previous work (Wang et al., 2018) and held constant across all architectures (see Table 3).

### Hidden unit analysis

Deep learning algorithms like RNNs are often described as “black boxes” due to difficulties in understanding the mechanisms underlying their behavior (Sussillo & Barak, 2013). To address this issue, hidden unit activity was analyzed in relation to behavior via dimensionality reduction techniques, in particular principal component analysis (PCA) and targeted dimensionality reduction (TDR, (Ebitz et al., 2018; Mante et al., 2013)). PCA was used to obtain a first intuition about internal dynamics, and TDR was used to analyze interpretable dimensions of the hidden unit data.

For PCA, hidden unit values were first centered and standardized. Then, the time course of the first three principal components (PCs) was plotted for individual RNN instances, and color-coded according to chosen option, state-value estimate, and stay versus switch decisions.

In contrast to PCA, TDR is a dimensionality reduction technique where the resulting high-dimensional neural activation data is projected onto axes with a specific interpretation. This is achieved by first using PCA to obtain an unbiased estimate of the most salient patterns of activations in the neural data and then regressing the resulting principal components against variables of interest. The resulting predicted values form the interpretable axes (Ebitz et al., 2018; Mante et al., 2013). Following previous work in primate neurophysiology (Ebitz et al., 2018; Mante et al., 2013), for discrete variables (e.g. choice, stay/switch behavior) we used logistic regression:

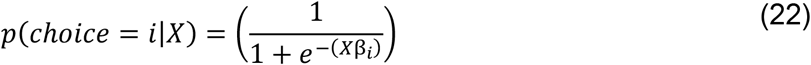

For continuous state-value estimates, we used linear regression:

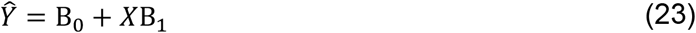

Here, X are the principal components based on the standardized hidden unit predictor matrix of size N (no. of trials) x M (no. of hidden units) and B_0_ *and* B_1_ are vectors of size M (no. of hidden units). The resulting axis 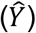, a vector of size no. of trials, now has a specific meaning - “Given the principal components on a given trial, what is the predicted value of the state-value-estimate?”. For discrete outcomes, the PCA-based hidden unit data were projected onto a choice predictive (or stay/switch-predictive) axis by inverting the logistic link function, i.e., for the case of the choice axis:

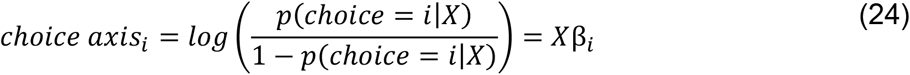

The *choice axis*_*i*_, a vector of size no. of trials, again has a concrete interpretation: “Given the de-noised hidden units on a trial, what are the log-odds of observing *choice*_*i*_”? If the log odds are positive, it is more likely, if it’s negative it is less likely to observe *choice*_*i*_. If predicted occurrence and non-occurrence of *choice*_*i*_ is equiprobable the log odds are 0.

To decode decisions from de-noised hidden unit activity, we used the results of the logistic regression (Equation 43) to calculate the probability of each action. The action with the maximum probability in a given trial was taken as the predicted action, and the proportion of correctly predicted choices was taken as the decoding accuracy.

---

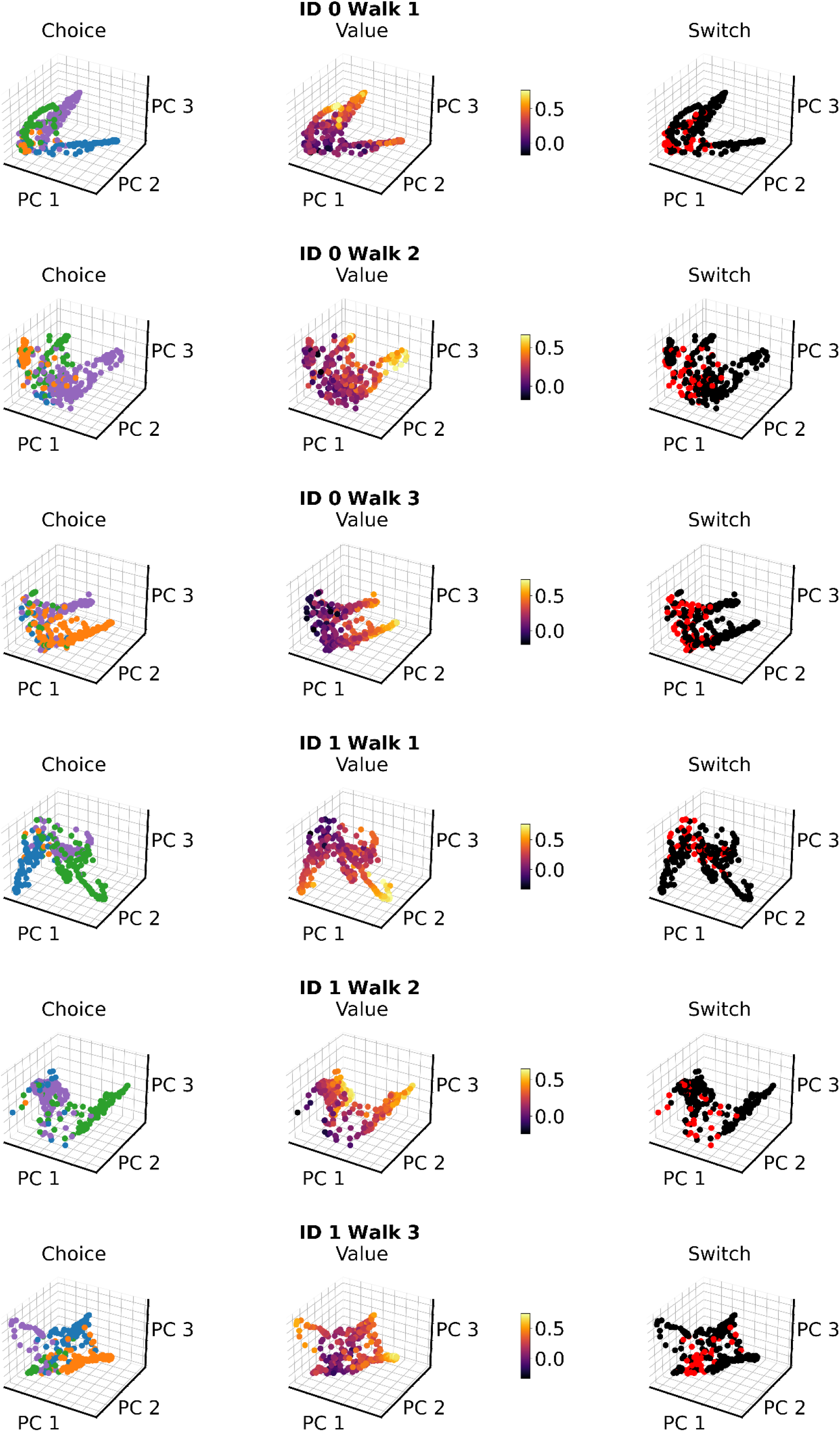

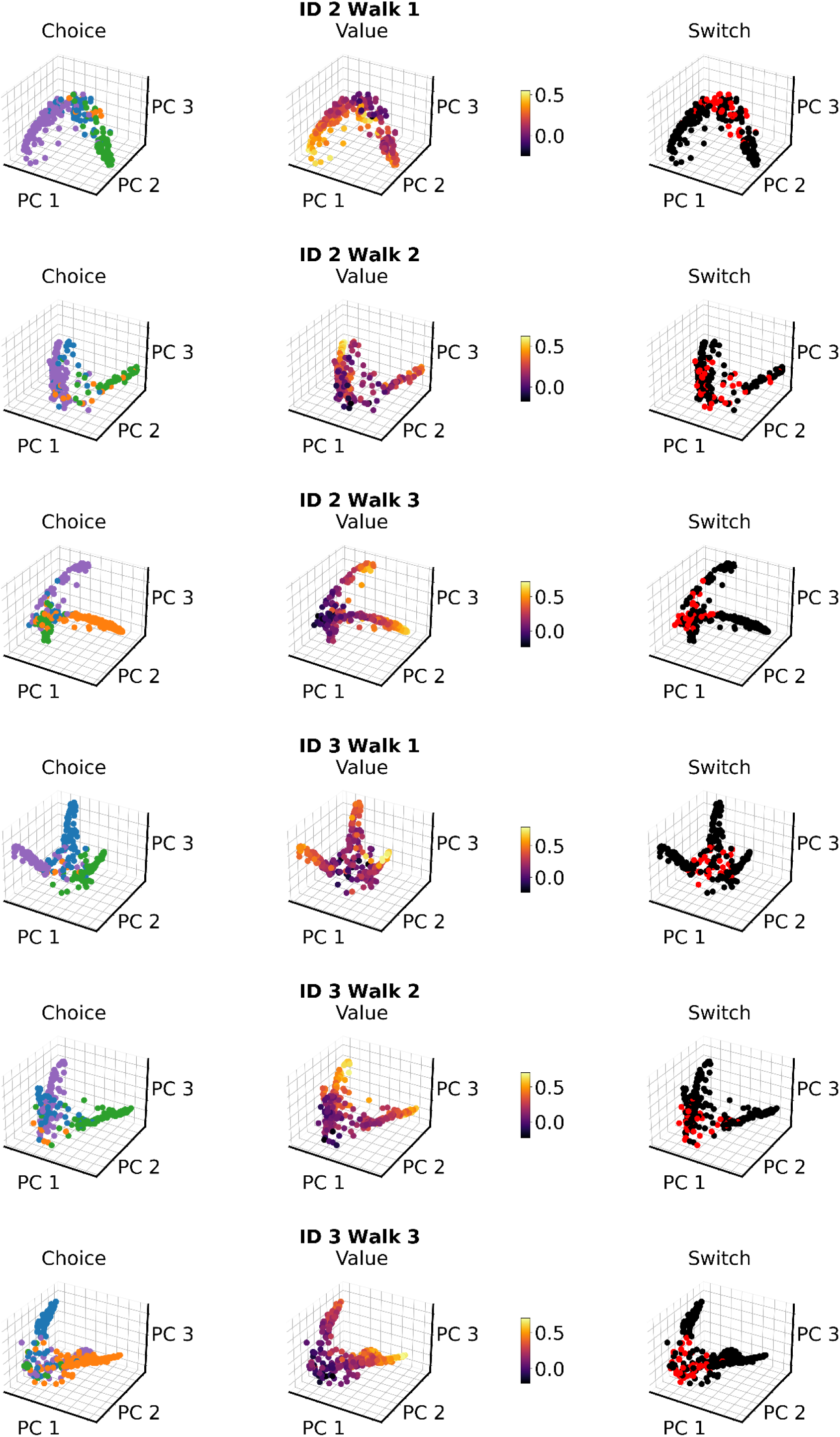

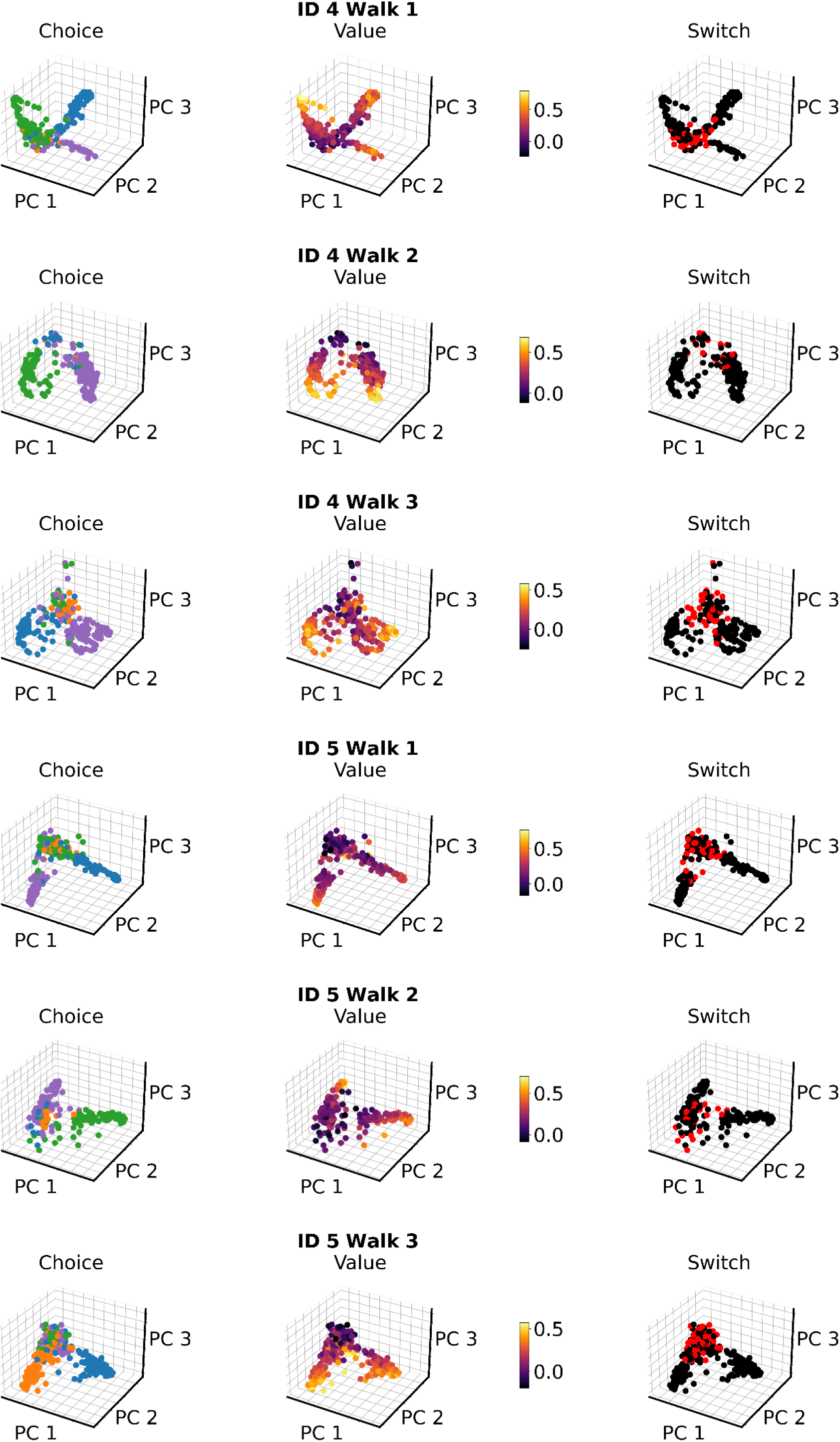

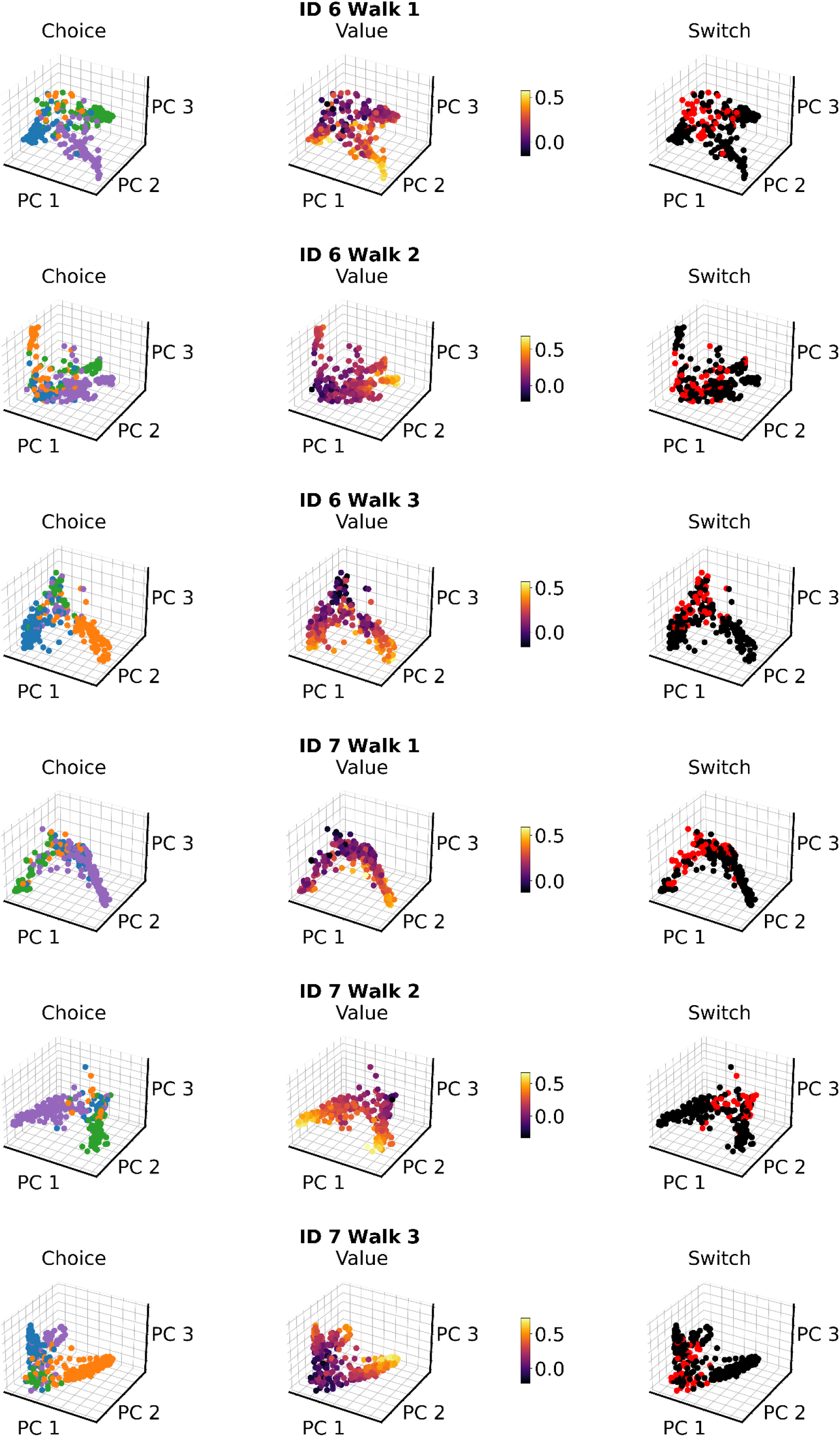

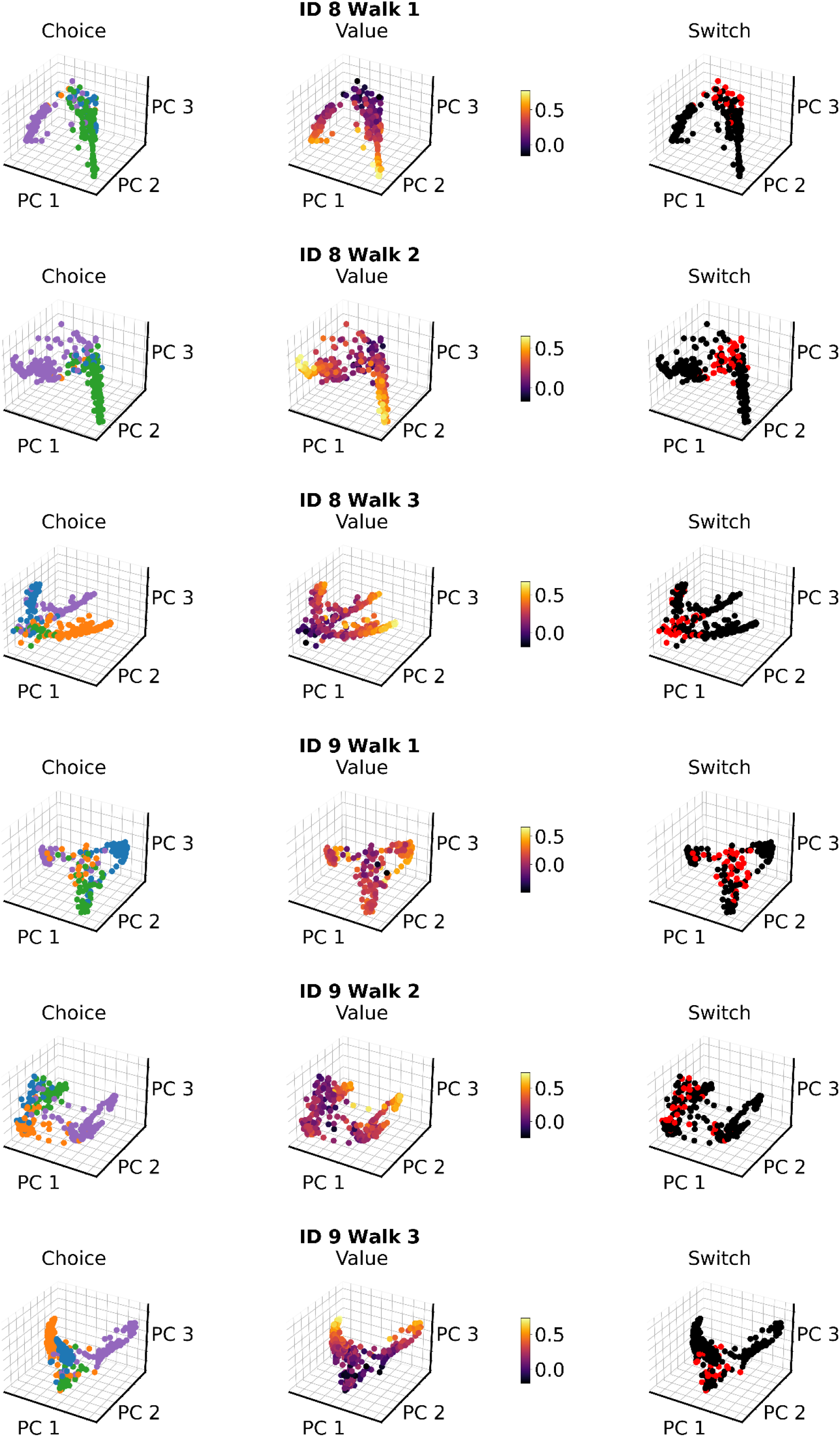

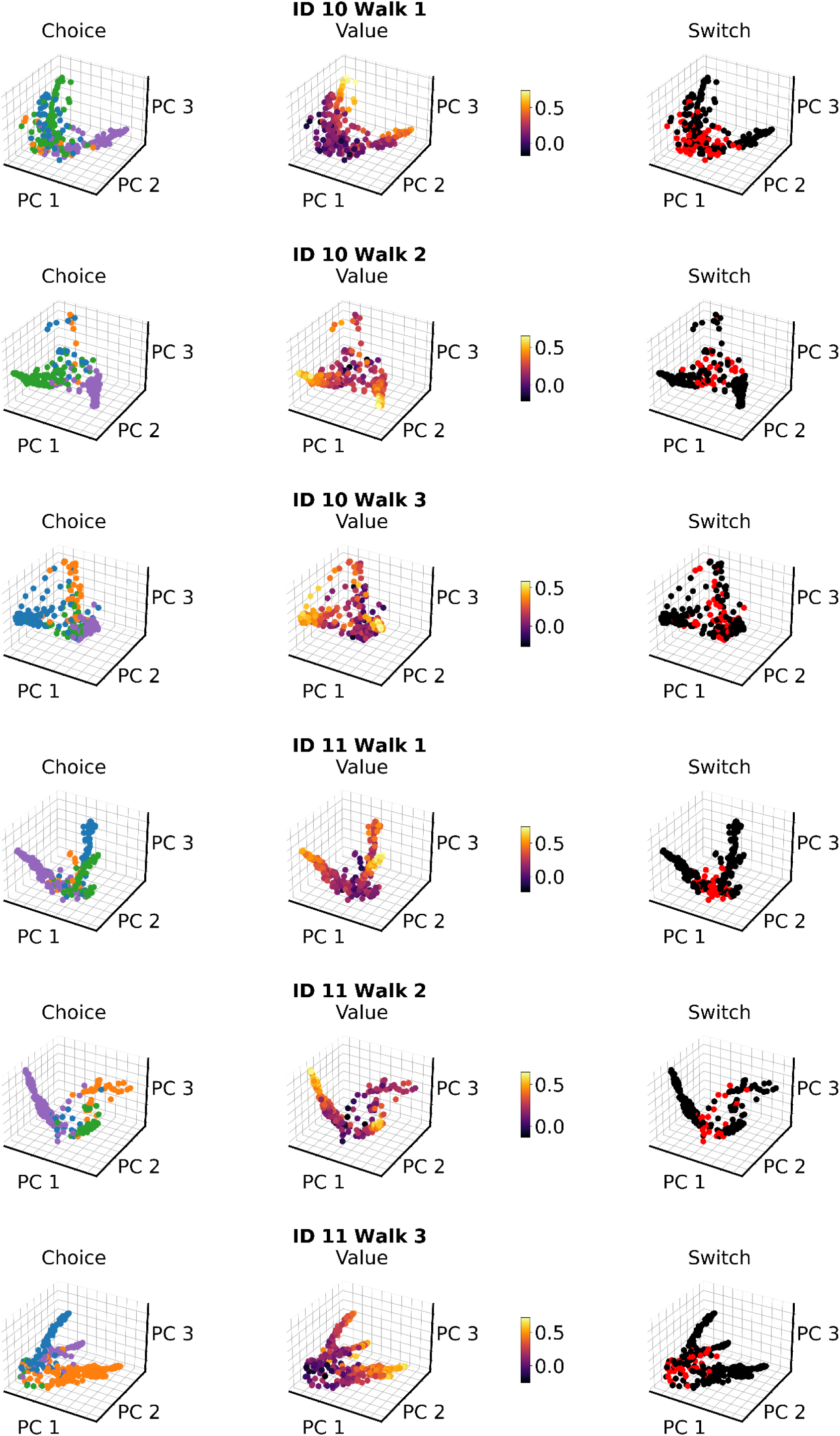

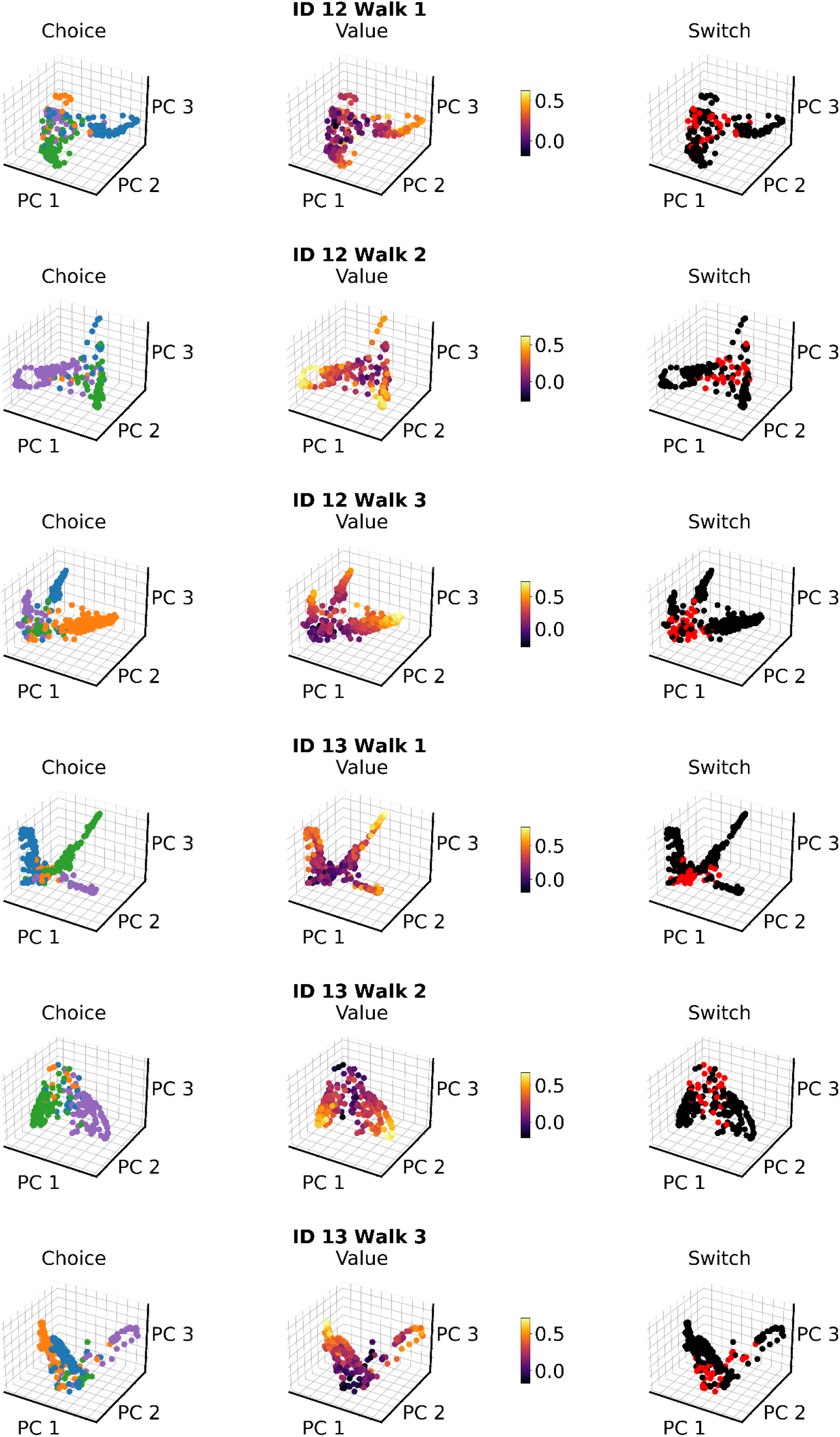

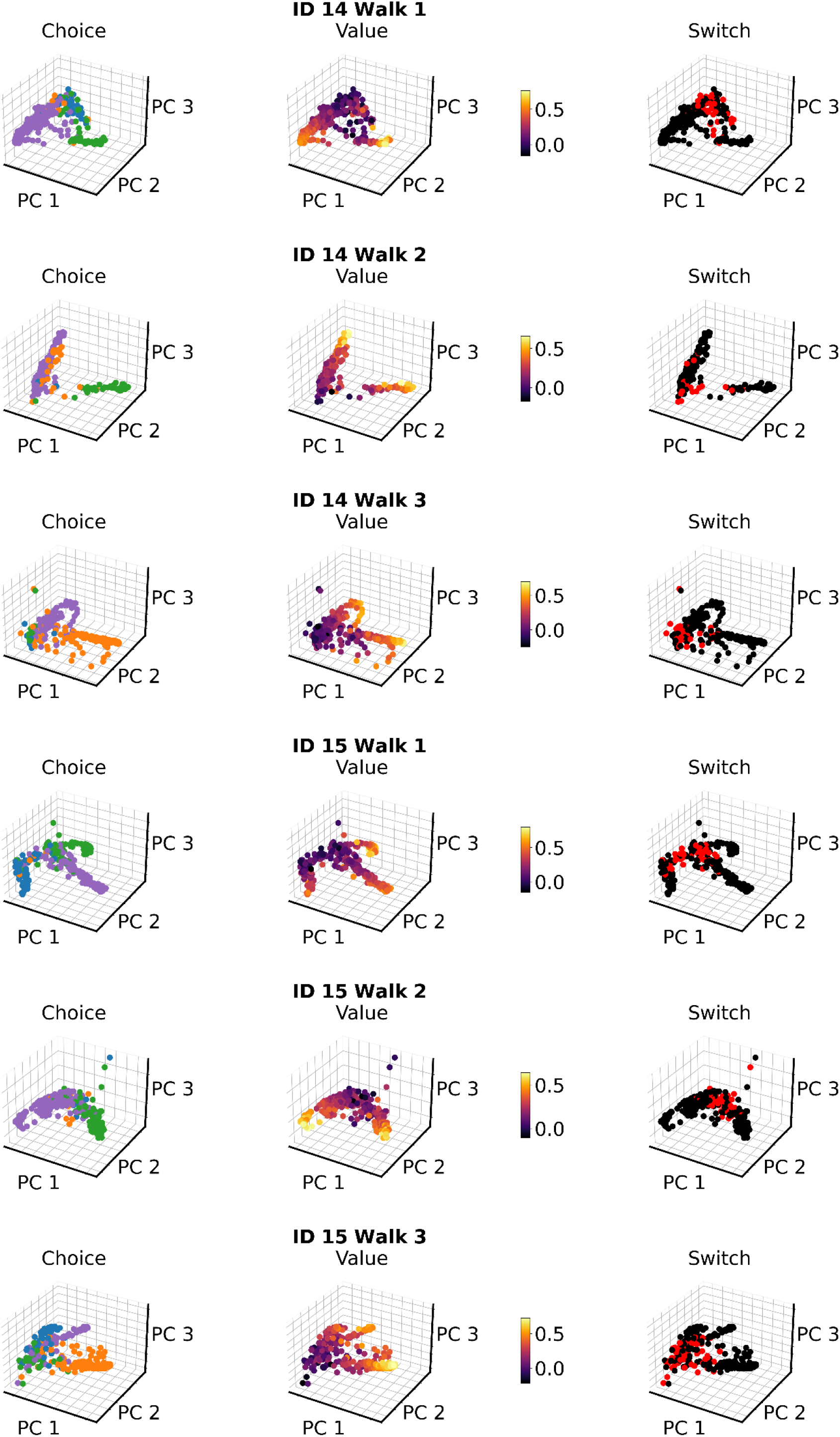

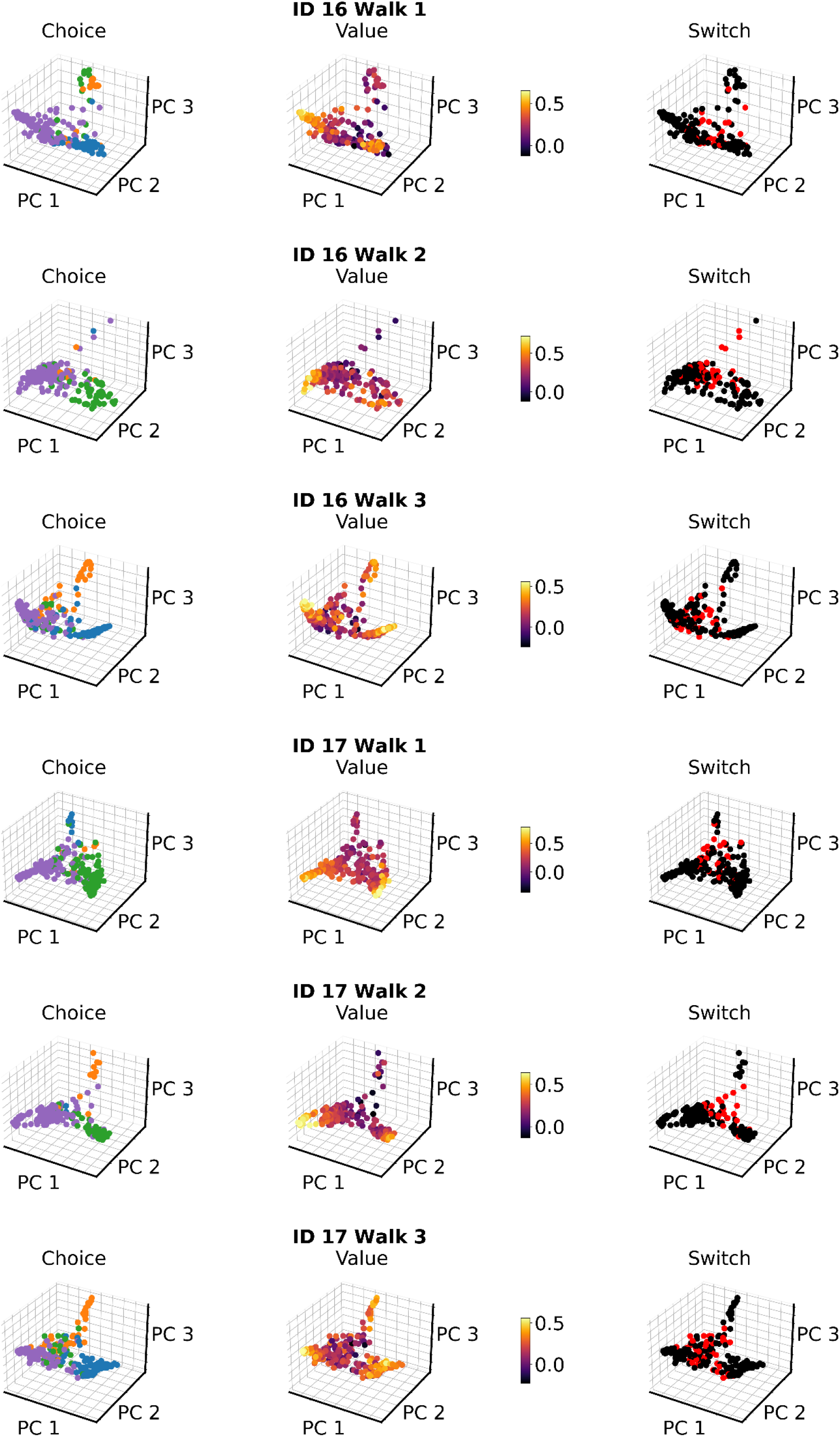

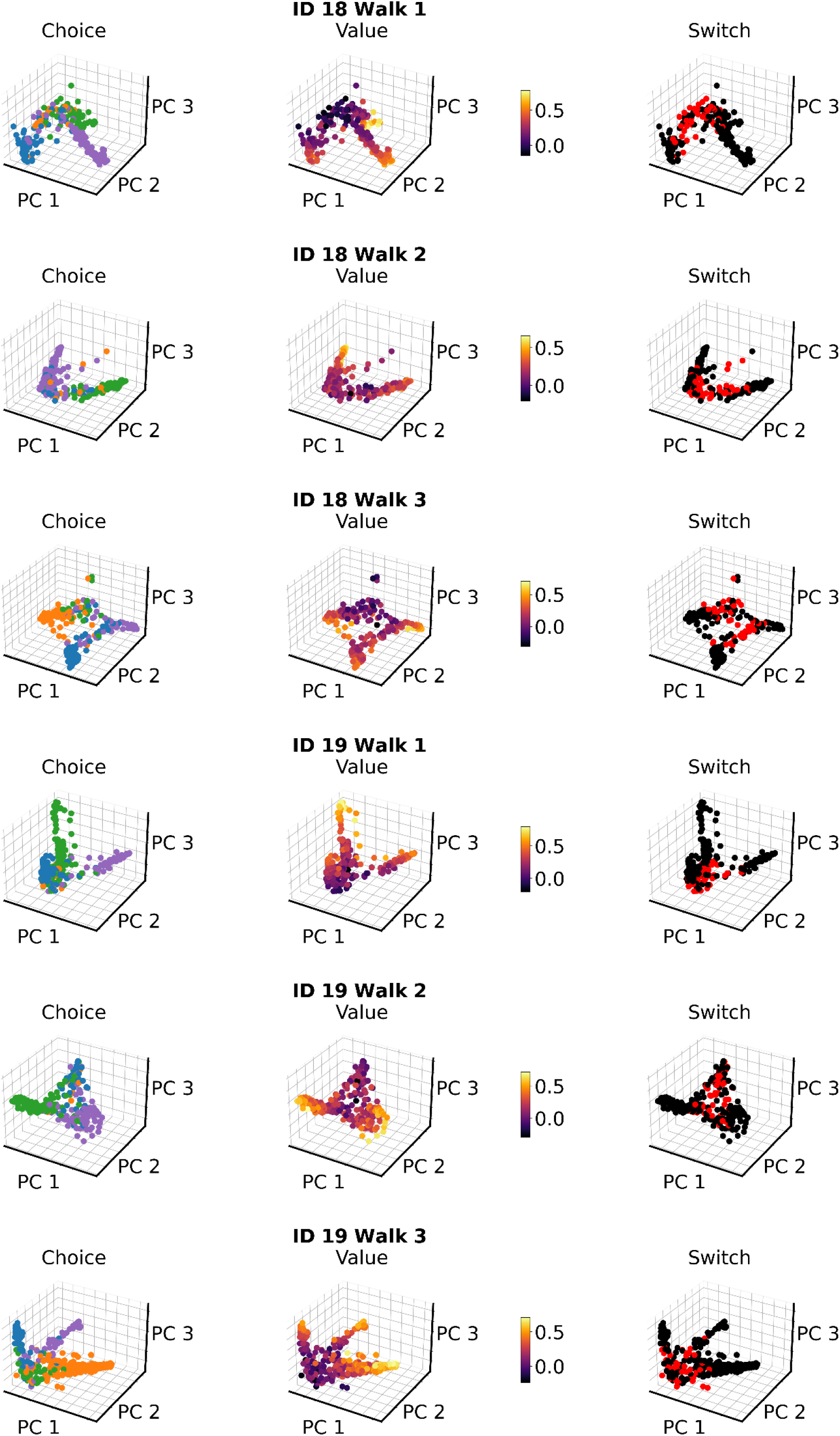

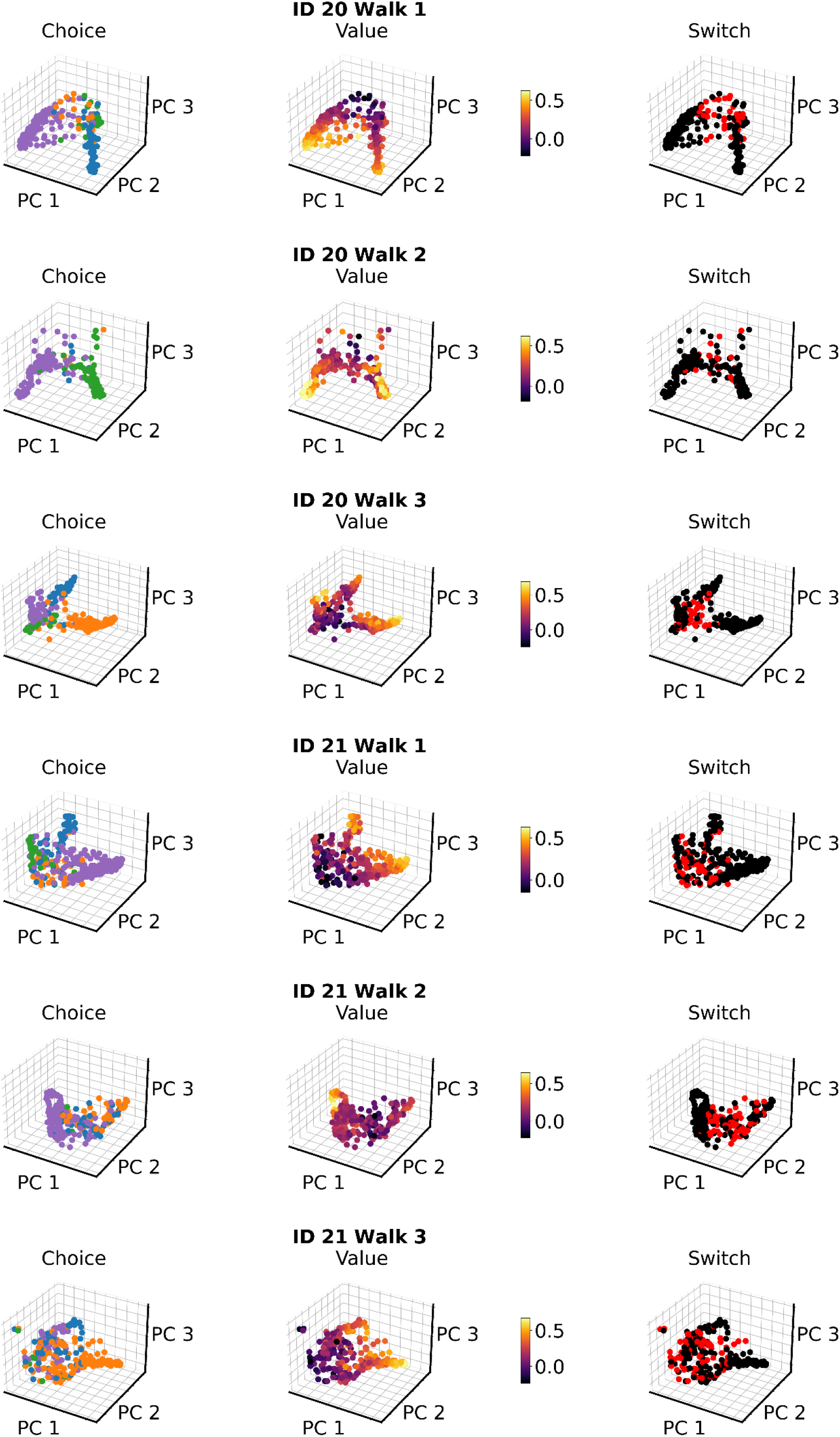

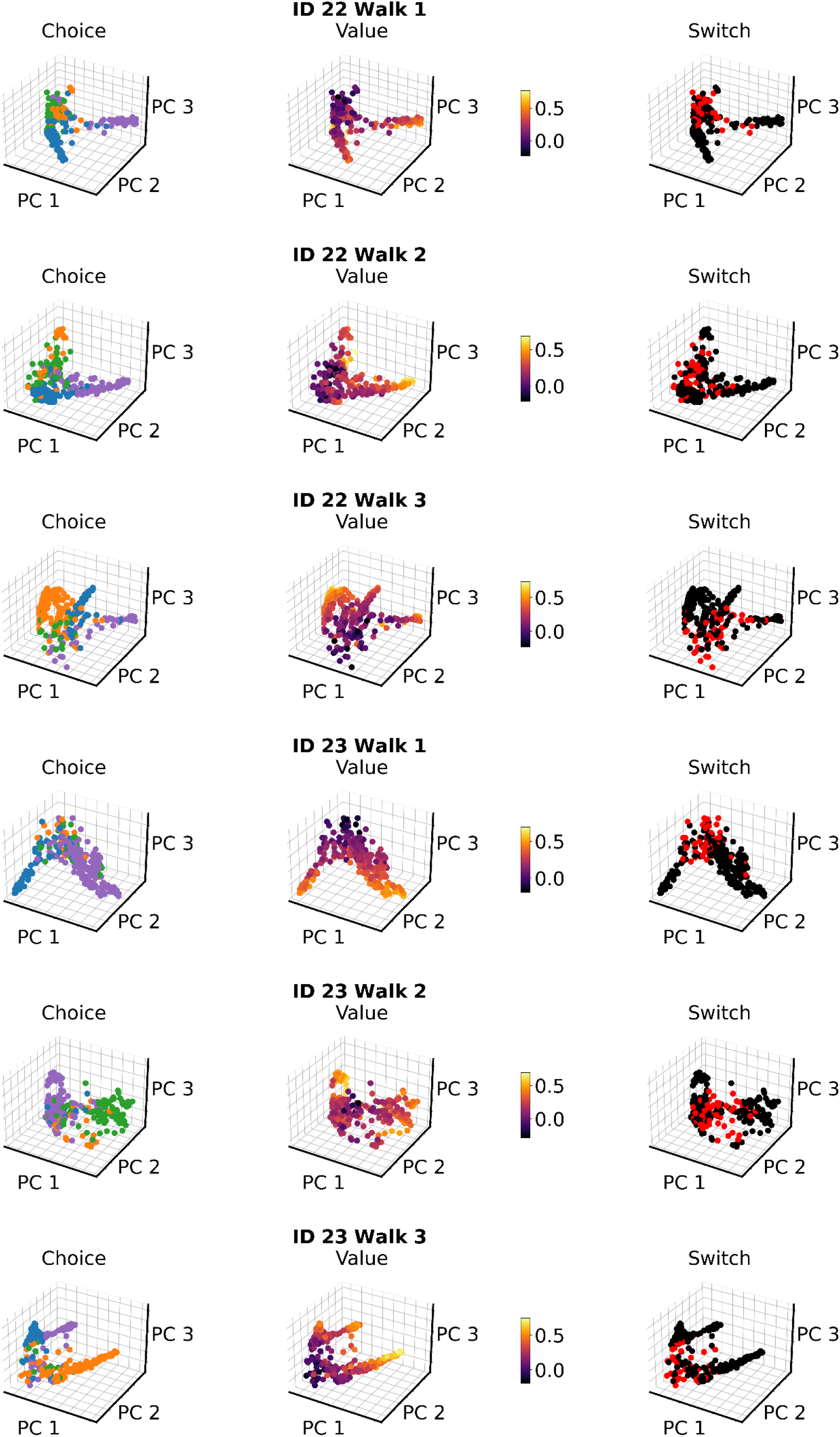

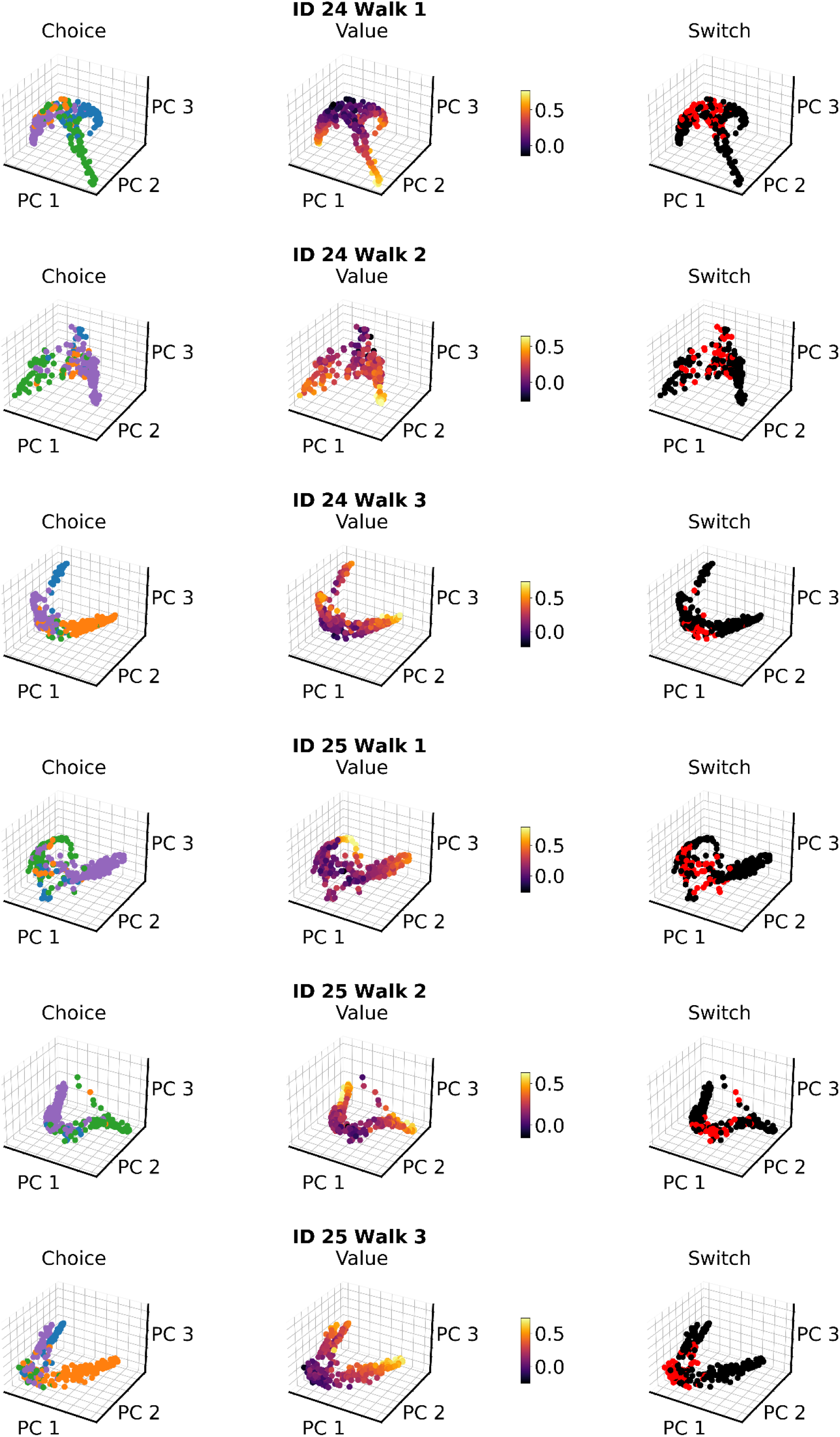

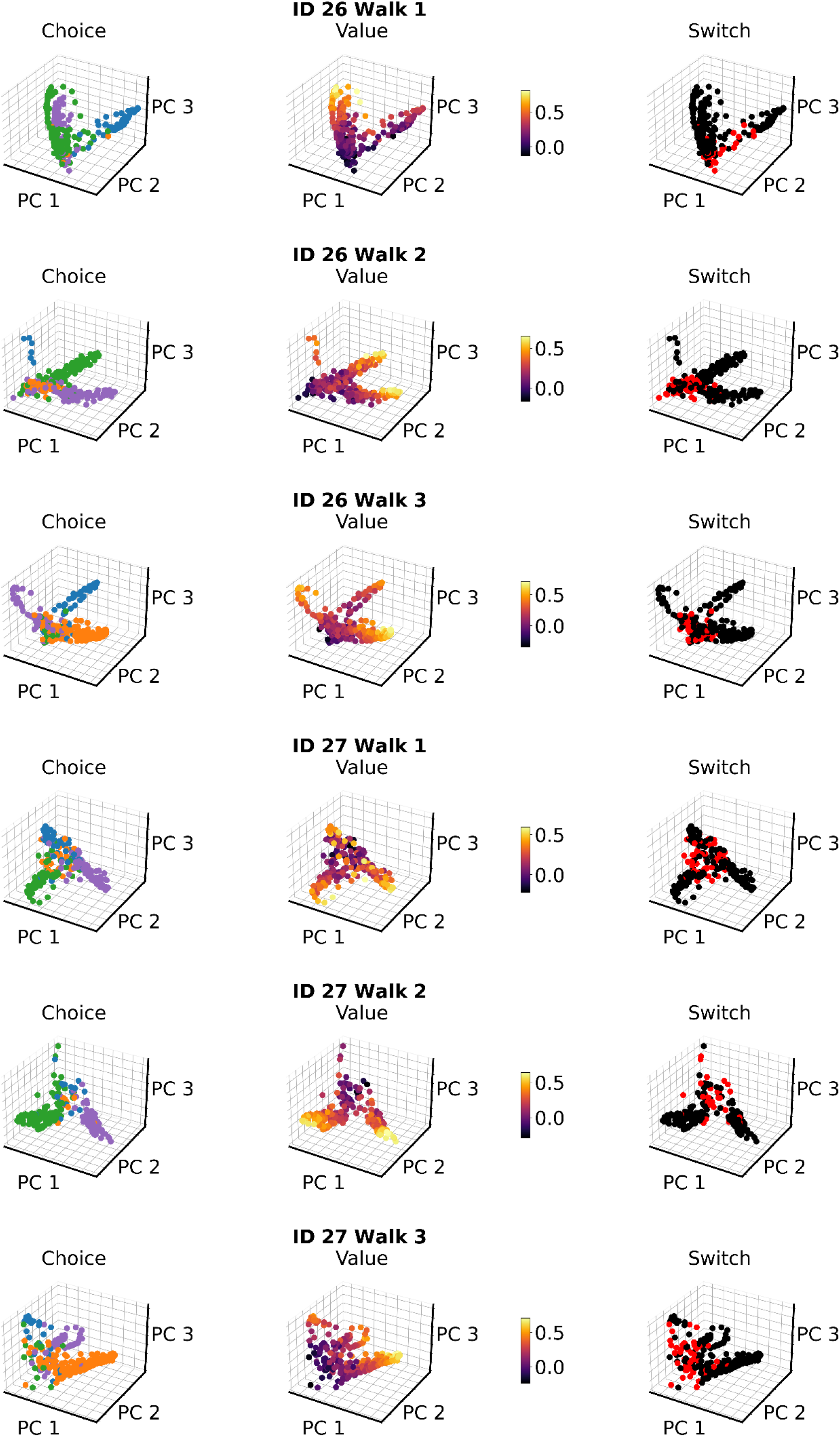

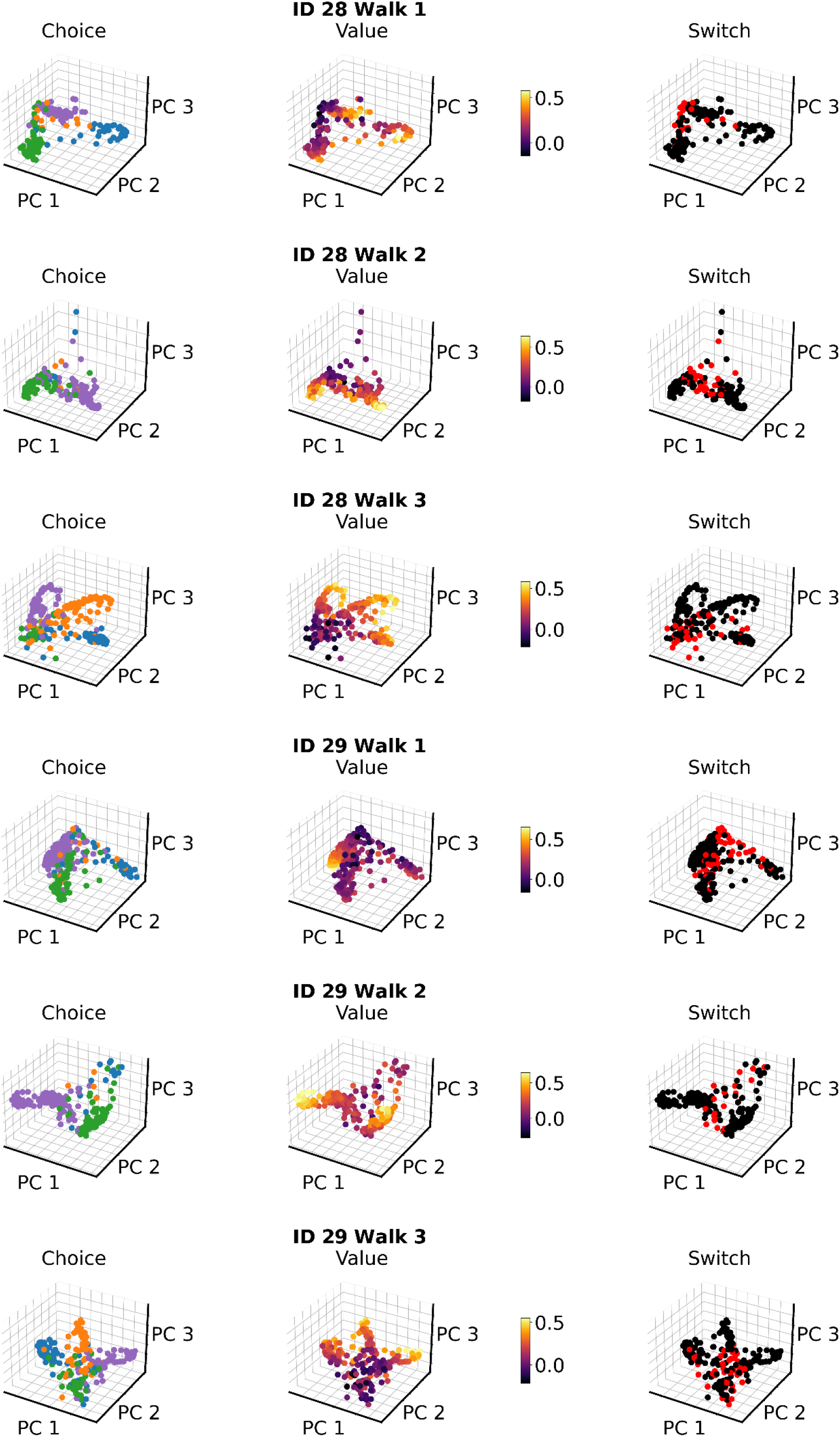

